# Non-enzymatic ABHD6 interacts with Akt-FoxO1 axis to regulate selective hepatic insulin resistance

**DOI:** 10.64898/2026.02.11.705361

**Authors:** Guannan Li, Laurence T. Maeyens, Jiyuan Yin, Jan-Bernd Funcke, Chanmin Joung, Ruizhen Li, Ziying Xu, Ting Wu, Xin Li, Nisi Jiang, Mbolle Ekane, Maria Paula Lopez, Pengju Cao, Sijia He, Adam B. Salmon, S.R. Murthy Madiraju, Marc Prentki, Juli Bai, James F. Nelson, Xianlin Han, Yi Zhu, Shangang Zhao

## Abstract

The enzymatic function of ABHD6 on insulin secretion and insulin resistance is well documented. However, its non-enzymatic function, especially its effects on selective hepatic insulin resistance and metabolic dysfunction-associated steatotic liver disease (MASLD) is completely unexplored. ABHD6 is elevated under conditions of diet-induced obesity and aging. To define the role of ABHD6 in liver physiology, we generated liver-specific ABHD6 knockout mice, as well as liver specific overexpression of native and enzymatic inactive mutant ABHD6 mouse models. We demonstrated that ABHD6 is an unidentified regulator of selective hepatic insulin resistance and contributes to MASLD and liver fibrosis. Furthermore, we found that non-enzymatic ABHD6, rather than its enzymatic form, contributes to this regulation. Mechanistically, we found that ABHD6 translocated into the nucleus and interacted with Akt/FoxO1 axis to regulate its function. In addition, knockdown of FoxO1 in primary hepatocytes or overexpression of constitutively active mutant FoxO1 by AAV approach could completely abolish the effects of ABHD6 on glucose tolerance and gluconeogenesis. Our study reveals an entirely different mechanism underlying selective hepatic insulin resistance that involves a previously unknown non-enzymatic function of ABHD6. This study opens an avenue for the development of a novel class of ABHD6 inhibitors to treat MASLD and liver fibrosis.

**Highlights:**

1. ABHD6 expression in the liver is increased with obesity and aging.
2. ABHD6 manipulation affects selective hepatic insulin resistance, MASLD and liver fibrosis.
3. Non-enzymatic ABHD6 interacts with Akt/FoxO1 axis to regulate FoxO1 transcriptional activity.
4. The effects of ABHD6 on glucose tolerance and hepatic gluconeogenesis are completely dependent on FoxO1 activity.

## Introduction

Selective hepatic insulin resistance is strongly linked with metabolic dysfunction-associated steatotic liver disease (MASLD), which has reached epidemic proportions worldwide^1,2^. In MASLD, insulin-mediated suppression of gluconeogenesis in the liver is impaired, while insulin-driven *de novo* lipogenesis appears to continue unimpeded^3^. Despite intensive research into the processes underlying selective insulin resistance in the liver, no definite picture has emerged to date. Both insulin signaling-related and - unrelated mechanisms have been proposed to contribute to selective hepatic insulin resistance^4,5^. These include relative decreases in glucose-handling compared to the lipid-handling arm of insulin signaling due to different intrinsic sensitivities of both arms and/or phosphorylation-dependent changes in Akt-substrate interactions^5–9^ as well as insulin signaling-independent increases in *de novo* lipogenesis due to extrahepatic factors such as free fatty acids, acetyl-CoA, glucose, fructose, and amino acids acting through substrate push, allosteric modulation, or transcriptional regulation^10–13^. However, even in their entirety, these mechanisms cannot fully explain the emergence of selective hepatic insulin resistance in MASLD, suggesting hitherto unrecognized contributors to this process to exist.

Akt (protein kinase B) serves as a central regulator of hepatic glucose and lipid metabolism downstream of insulin signaling^3,14^. Once insulin binds to its receptor, a signaling cascade activates Akt to promote liver *de novo* lipogenesis by modulating downstream targets such as mTORC1, Insig2a, and ACLY^8,15–17^. At the same time, Akt regulates liver glucose metabolism by inhibiting glycogen synthase kinase 3 (GSK3α/β), resulting in increased glycogen synthase activity, and by inhibiting forkhead box O family members, including FoxO1 and FoxO3a, resulting in decreased gluconeogenesis^14,18–20^. Given the dual function of Akt in hepatic lipid and glucose metabolism, Akt has been proposed as a critical bifurcation point in insulin signaling, especially in the context of selective hepatic insulin resistance^21^. Despite broad agreement on diminished Akt responsiveness and activity under insulin-resistant conditions, it’s role in selective hepatic insulin resistance remains poorly understood^22^. Emerging evidence suggests that Akt may exhibit substrate selectivity^23,24^; however, whether this selectivity contributes to the development of selective hepatic insulin resistance has not been investigated.

α/β hydrolase domain-containing 6 (ABHD6) is a lipase that has been reported to hydrolyze monoacylglycerol (MAG), bis(monoacylglycero)phosphate (BMP), and lysophosphatidylserine (LysoPS)^25–28^. ABHD6 is broadly expressed throughout the body and has been implicated in multiple pathophysiological progresses, including obesity, insulin resistance, diabetes, Alzheimer’s disease, and multiple sclerosis^29,30^. We and others previously demonstrated that inhibition of ABHD6 promotes glucose-induced insulin secretion from pancreatic islets and enhances the thermogenic activity of adipose tissue^30–33^. However, its role in MASLD, especially in selective hepatic insulin resistance, has not been explored. In this study, we reveal a crucial function of non-enzymatic ABHD6 as a driver of selective hepatic insulin resistance and regulator of MASLD progression by modulating the Akt-FoxO axis.

## Results

### Liver ABHD6 expression is increased with obesity and aging

To study the role of ABHD6 in MASLD, we first examined its expression in the liver in the settings of obesity and aging, two conditions known to promote the development of MASLD. Of note, we observed no differences in liver ABHD6 protein or *Abhd6* mRNA expression comparing chow-fed male and female wildtype mice (Fig. 1A-B) and thus decided to perform all our subsequent experiments in male mice. Feeding male wildtype mice with a high-fat diet (HFD), resulted not only in liver insulin resistance (Sup Fig. 1A) but also a 2-fold increase in ABHD6 protein and *Abhd6* mRNA expression (Fig. 1C-D). Comparing old to young male wildtype mice, we detected a robust increase in the expression of the senescence markers p53 and p16 as well as a robust 2-fold increase in ABHD6 protein expression, while no changes in *Abhd6* mRNA expression were observed (Fig. 1E-F). Taken together, these results establish that obesity and aging are accompanied by a substantial increase of ABHD6 expression in the liver, suggesting a potential role of ABHD6 in MASLD development.

**Figure 1:**
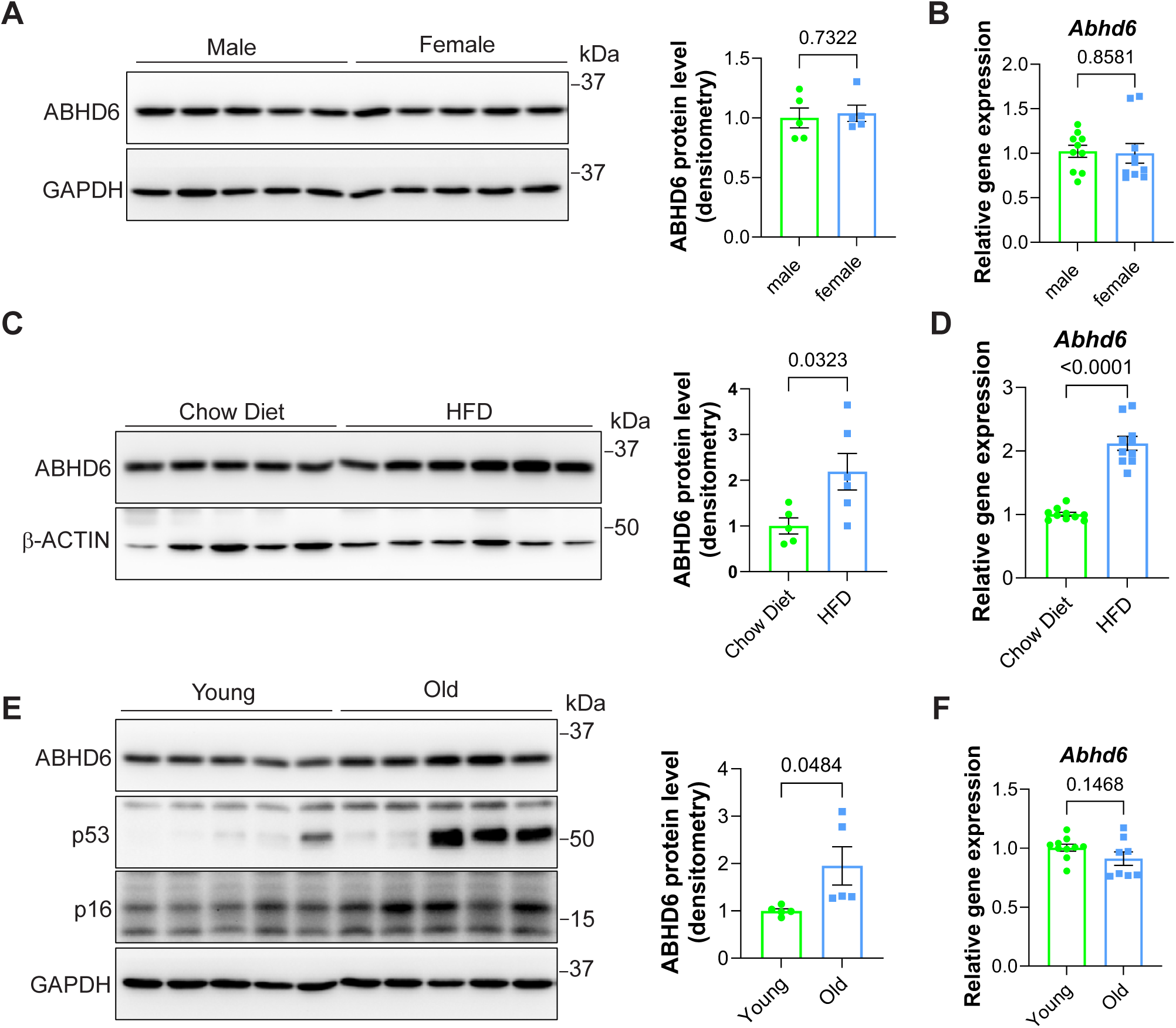
Liver ABHD6 expression is increased with obesity and aging. **(A and B)** 8-week-old male and female wildtype mice on a chow diet were compared (n = 5 per group). **(A)** ABHD6 protein levels in the liver. **(B)** *Abhd6* mRNA levels in the liver (n = 10 per group). **(C and D)** 8-week-old male wildtype mice were fed a chow diet or high-fat diet (HFD) for 8 weeks (n = 5-6 per group). **(C)** ABHD6 protein levels in the liver. **(D)** *Abhd6* mRNA levels in the liver (n =10 per group). **(E and F)** 6-month-old (‘young’) and 16-month-old (‘old’) male wildtype mice on a chow diet were compared (n = 5 per group). **(E)** ABHD6, p53, and p16 protein levels in the liver. **(F)** *Abhd6* mRNA levels in the liver (n = 10 per group). (A-F) Bar graph data are displayed as mean ± SEM and were analyzed by Student’s *t* test. p-values of respective comparisons are provided.

Hepatocyte-specific deletion of ABHD6 alleviates the metabolic impact of MASLD and impedes its progression to MASH.

As liver ABHD6 expression is significantly increased in diet-induced MASLD, we wondered whether reducing ABHD6 expression in hepatocytes could prevent the development of MASLD. To this end, we generated a hepatocyte-specific ABHD6 knockout model (LABHD6 KO) by crossing *Albumin-Cre* with *Abhd6 floxed* mice. Immunoblot and qPCR analyses demonstrated a significant reduction in liver ABHD6 protein and *Abhd6* mRNA expression in LABHD6 KO mice compared to littermate control mice (Fig. 2A-B). Importantly, no changes were observed in ABHD6 protein expression in the epididymal white adipose tissue (eWAT) of LABHD6 KO mice, confirming the tissue-specificity of our knockout approach (Fig. S2A). First, we addressed whether deletion of ABHD6 would affect liver function under chow fed conditions. We found that LABHD6 KO mice exhibited comparable body weight gain and glucose tolerance to their littermate control mice (Fig. S2B-C), indicating that deletion of ABHD6 does not affect overall energy homeostasis under chow fed conditions.

**Figure 2:**
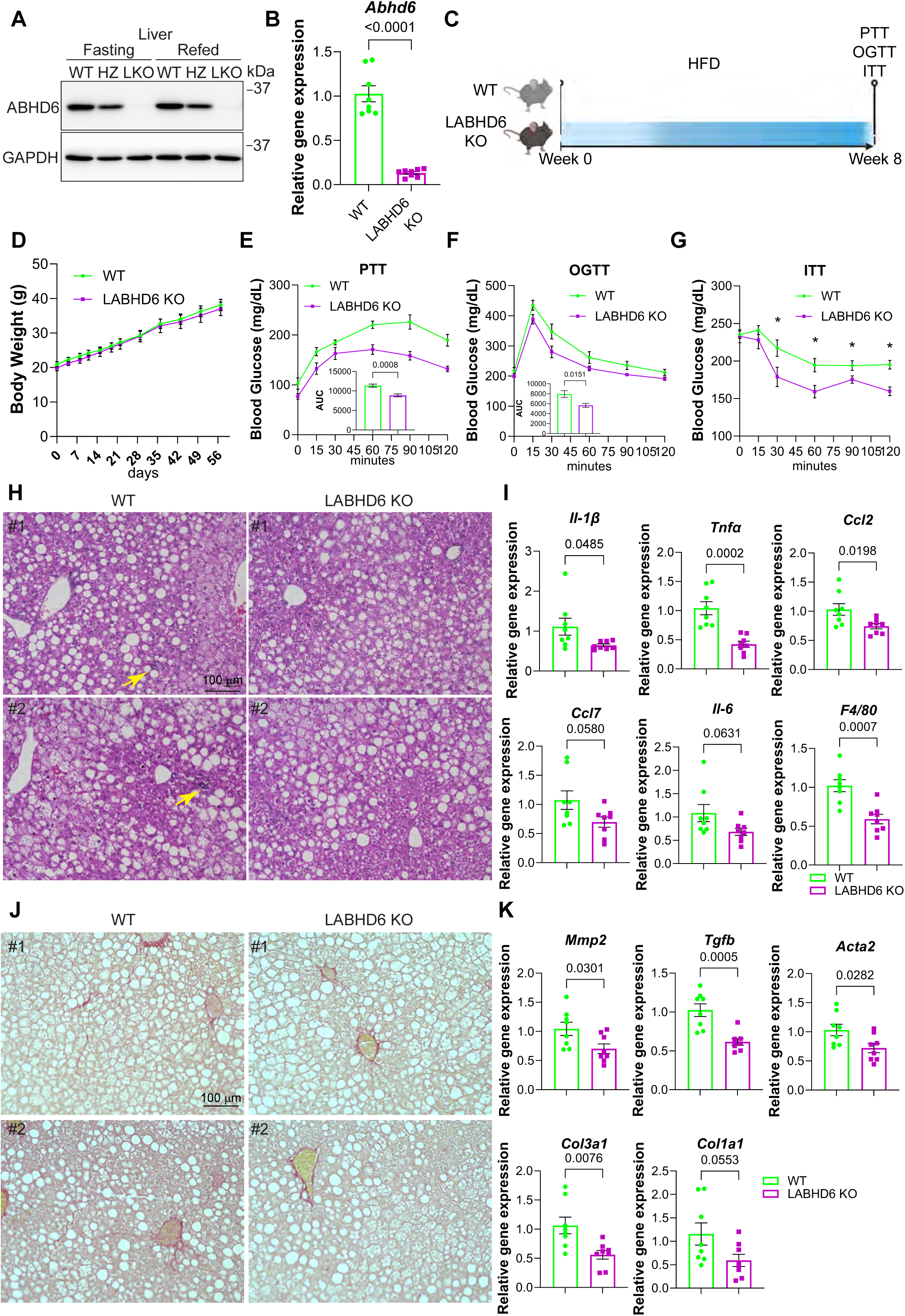
Hepatocyte-specific deletion of ABHD6 alleviates the metabolic impact of MASLD and impedes its progression to MASH. **(A)** 8-week-old male littermate control (WT) mice, heterozygous LABHD6 KO (HZ) mice, and homozygous LABHD6 KO (LKO) mice were compared. ABHD6 protein levels in the livers were analyzed. **(B)** *Abhd6* mRNA levels in the livers (n = 8 per group). **(C-K)** 8-week-old littermate control (WT) and homozygous LABHD6 KO mice were placed on high-fat diet (HFD) for 8 weeks. **(C)** Overview of experimental setup. **(D)** Body weight (n = 7 per group). **(E-G)** Blood glucose levels during pyruvate tolerance test (PTT), glucose tolerance test (GTT), and insulin tolerance test (ITT). For PTT, mice were starved O/N, and for OGTT and ITT, mice were deprived of food access for 6 hours. For the PTT and GTT, the area under the curve (AUC) was calculated (n = 7 per group). **(H)** H&E staining of livers (n = 2 per group). Arrows indicated clusters of immune cells. **(I)** mRNA expression of inflammation markers (n = 8 per group). **(J)** Picrosirius red staining of livers (n = 2 per group). **(K)** mRNA expression of fibrosis markers (n = 8 per group). (B, D-G, I, K) Data are displayed as mean ± SEM and were analyzed by Student’s *t* test (bar graphs) or one-way ANOVA (curves). p-values of respective comparisons are provided. *p < 0.05.

Next, we challenged LABHD6 KO and littermate control mice with a HFD and assessed MASLD progression (Fig. 2C). ABHD6 elimination had no effect on body weight or liver, inguinal white adipose tissue (iWAT), and eWAT weight (Fig. 2D and S2D). Strikingly though, LABHD6 KO mice performed significantly better in pyruvate, glucose, as well as insulin tolerance tests, indicating improved metabolic control (Fig. 2E-G). Skeletal muscle and adipose tissue are major organs regulating blood glucose homeostasis. We therefore examined insulin signaling in skeletal muscle from LABHD6 knockout mice. Neither ABHD6 protein levels nor insulin signaling were altered in muscle (Fig. S2E), suggesting that hepatic ABHD6 is likely the primary contributor to blood glucose regulation in LABHD6 knockout mice. Interestingly, H&E staining revealed comparable lipid accumulation in the livers of LABHD6 KO and littermate control mice (Fig. 2H). The deletion of ABHD6 did however greatly reduce signs of inflammation in the liver, such as the expression of *Il1b*, *Tnf*, *Ccl2*, *Ccl7*, *Il6*, and *Adgre* (F4/80), *Il1a*, and *Cxcl10* mRNA (Fig. 2H-I and S2F), suggesting slowed progression from MASLD to metabolic dysfunction-associated steatohepatitis (MASH). The occurrence of fibrotic changes in the liver is another crucial indicator of disease progression. We thus also examined the manifestation of liver fibrosis in LABHD6 KO mice. Doing so, we found ABHD6 elimination to greatly reduce signs of fibrosis in the liver, reflected by decreased collagen accumulation (picrosirius red staining; Fig. 2J) and reduced expression of *Mmp2*, *Tgfb1*, *Acta2* (*Sma*), *Col3a1*, and *Col1a1* mRNA (Fig. 2K). Taken together, these results clearly indicate that eliminating ABHD6 from hepatocytes alleviates the systemic metabolic impact of diet-induced MASLD and diminishes liver inflammation and fibrosis.

Hepatocyte-specific overexpression of ABHD6 in liver exacerbates the metabolic impact of MASLD and promotes its progression to MASH.

To corroborate our findings, we also generated a hepatocyte-specific ABHD6 overexpression model (LABHD6 OE) *via* tail vein injection of AAV8 TBG-*Abhd6* into male wildtype mice, with AAV8 TBG-eGFP serving as control. One-week post-injection, we placed LABHD6 OE and control mice on an HFD for 12 weeks (Fig. 3A). We first confirmed the specificity of ABHD6 overexpression by analyzing liver and eWAT by immunoblot and RT-qPCR. These analyses demonstrated that *Abhd6* mRNA expression and ABHD6 protein were significantly increased in liver, but unchanged in eWAT (Fig. 3B-C and S3A-B), validating our viral overexpression approach. ABHD6 overexpression had no effect on body weight or liver, iWAT, and eWAT weight (Fig. 3D-E). Complementing our observations in LABHD6 KO mice, LABHD6 OE performed significantly worse in pyruvate, glucose, as well as insulin tolerance tests (Figure 3F-H), indicating deteriorated metabolic control. In addition, we found that overexpression of ABHD6 in hepatocytes significantly increased liver inflammation, despite reduced lipid droplet accumulation (Fig. 3I-J). This increase in liver inflammation was reflected by elevated *Il1b*, *Tnf*, *Ccl2*, *Ccl7*, *Il6*, and *Adgre* (*F4/80*) mRNA levels in LABHD6 OE mice (Figure 3J). Accompanying increased inflammation, we also LABHD6 OE mice also developed liver fibrosis, reflected by increased collagen accumulation (picrosirius red staining; Fig. 3K) as well as increased *Tgfb1* and *Col3a1* mRNA expression (Fig. 3L). In summary, these results strongly support the notion of hepatocyte ABHD6 as a hitherto underappreciated contributor to the systemic metabolic deterioration in MASLD and its progression to MASH.

**Figure 3:**
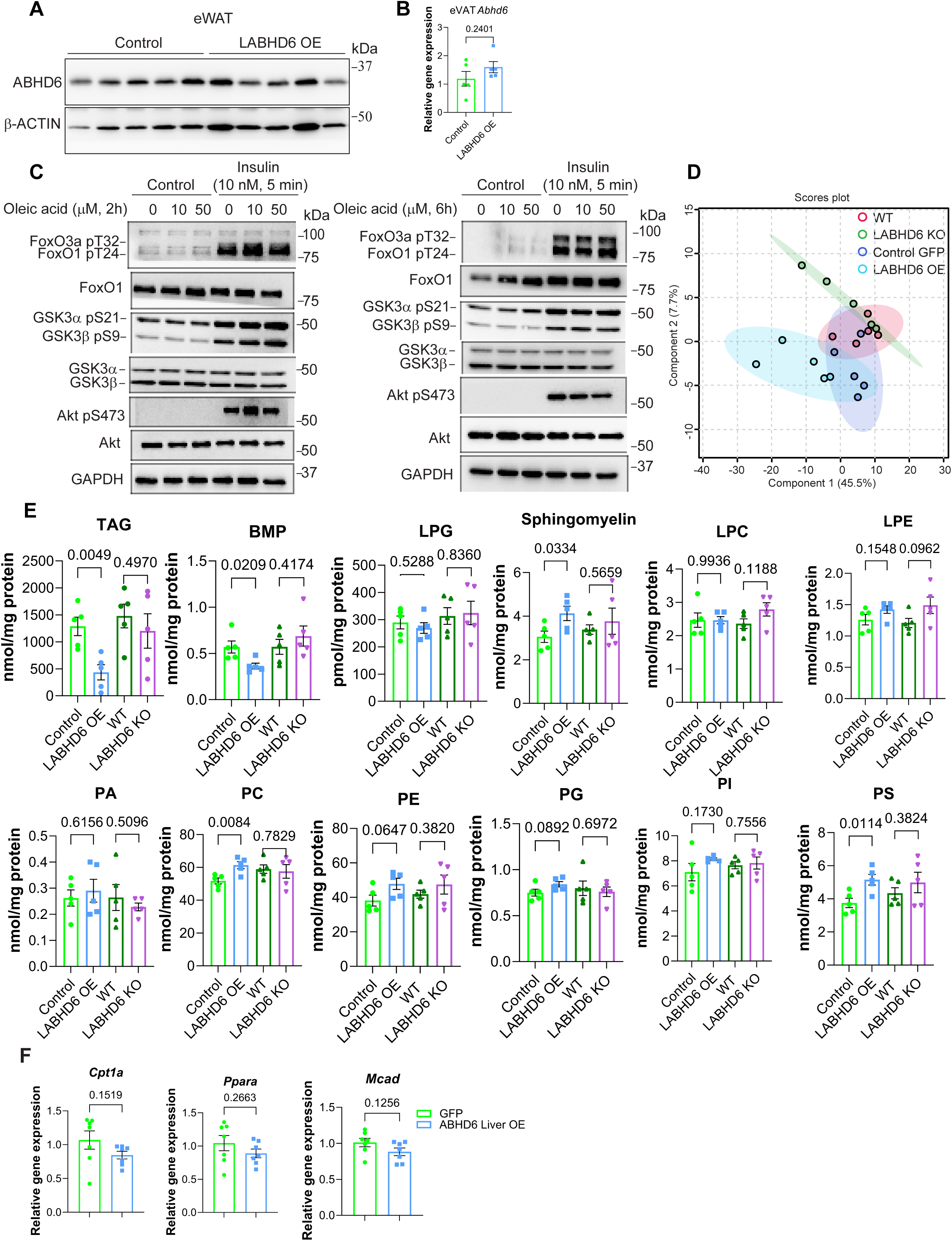
Hepatocyte-specific overexpression of ABHD6 in liver exacerbates the metabolic impact of MASLD and promotes its progression to MASH. 8-week-old male wildtype mice were injected with 1×10¹¹ genome copies AAV8 TBG-eGFP (Control), TBG-*Abhd6* (LABHD6 OE) or TBG-*Abhd6* p.S148A (LABHD6 S148A OE). 1-week post-injection, the mice were fed a high-fat diet (HFD) for 12 weeks. **(A)** Overview of experimental setup. **(B)** *Abhd6* mRNA levels in the liver (n = 10 per group). **(C)**ABHD6 protein levels in the liver (n = 5 per group). **(D and E)** Body weight and organ weights (n = 7 per group). **(F-H)** Blood glucose levels during pyruvate tolerance test (PTT), glucose tolerance test (GTT), and insulin tolerance test (ITT). For the PTT and GTT, the area under the curve (AUC) was calculated (n=6 per group). **(I)** H&E staining of livers (n = 2 per group). Arrows indicate clusters of immune cells. **(J)** mRNA expression of inflammation markers (n = 7 per group). **(K)** Picrosirius red staining of livers (n = 2 per group). **(L)** mRNA expression of fibrosis markers (n=7 per group). **(M)** Blood glucose levels during pyruvate tolerance test (PTT. For the PTT, AUC was calculated (n=7 per group). **(N)** Body weight (n = 7 per group). (B, D-H, J, L-N) Data are displayed as mean ± SEM and were analyzed by Student’s *t* test (bar graphs) or one-way ANOVA (curves) p-values of respective comparisons are provided. *p < 0.05.

### ABHD6-accessible lipids do not mediate the effects of ABHD6 on insulin resistance

As ABHD6 hydrolyzes monoacylglyerol (MAG) in multiple tissues, we tested whether elevating 1-MAG levels might mediate the effects of ABHD6 on the insulin signaling cascade. To this end, we treated mouse primary hepatocytes with 1-oloylglycerol (1-OG) at two different doses (10 µM or 50 µM) and time spans (2 hours and 6 hours). Independent of the dose or time span, we observed no effect of 1-OG treatment on insulin signaling (Fig. S3C), strongly arguing against an involvement of 1-MAG in the effects of ABHD6 on insulin resistance.

Recent studies have indicated that ABHD6 not only hydrolyzes MAG but also bis(monoacylglycero)phosphate (BMP), and lysophosphatidylserine (LysoPS)^25–28^. We thus investigated whether overexpressing or eliminating ABHD6 in hepatocytes could induce a change in the lipid profile, which may in turn affect hepatic insulin sensitivity. To this end, we performed a comprehensive lipidomic analysis in the livers of LABHD6 KO and OE mice. Processed data was clustered in partial least squares discriminant analysis (PLS-DA) for both cohorts. Based on absolute lipid abundances, the overall lipidome showed moderate separation of the WT-LABHD6 KO groups and the GFP-LABHD6 OE groups. (Fig. S3D). Similarly, we did not observe any significant and reciprocal changes in total lipid species in those four mouse models, including triacylglycerol (TAG), BMP, phosphatidylglycerol (PG), lysoPG, phosphatidylcholine (PC), lysoPC, phosphatidylethanolamine (PE), lysoPE, phosphatidic acid (PA), phosphatidylserine (PS), phosphatidylinositol (PI) and sphingomyelin (Fig. S3E and Table S2). Total TAG level was significantly reduced only in the livers of LABHD6 OE mice (Fig. S3E), consistent with the H&E staining results (Fig. 2H and 3I), despite unchanged lipogenesis and fatty acid oxidation (Fig. S6E and S3F). Similarly, total lipid species including BMP, PE, PC, PS and sphingomyelin were altered only in the livers of LABHD6 OE mice (Fig. S3E). Among all the lipid species that were significantly and reciprocally changed in the livers of LABHD6 KO and OE mice, we found that only one specific BMP species, namely BMP 20:4-22:6, was increased in LABHD6 KO and decreased in LABHD6 OE mice (Fig. S4A-C). Carefully comparing all BMP classes, we found that elimination of ABHD6 resulted in a trend towards increased levels of most BMP classes, whereas overexpression of ABHD6 lead to drastically decreased levels of most BMP classes (Fig. S4D). These results provide additional evidence that ABHD6 is a major hydrolase for BMP.

As BMP is the only lipid species that changed in response to both eliminating and overexpressing ABHD6, we investigated whether BMP serves as the mechanistic link between ABHD6 and insulin resistance. We thus treated mouse primary hepatocytes with BMP for two different time spans (2 hours and 16 hours). Doing so, we found that exogenous BMP did not affect the insulin signaling cascade (Fig. S4E). Collectively, these findings led us to explore a lipid-independent mechanism by which ABHD6 modulates insulin resistance.

To this end, we generated AAV8-TBG-*Abhd6* p.S148A, which expresses an enzymatically inactive ABHD6 mutant^25^. We then established a liver-specific ABHD6 S148A overexpression (LABHD6 S148A OE) mouse model by tail-vein injection of AAV8-TBG-*Abhd6* p.S148A into male wild-type mice, using AAV8-TBG-eGFP and AAV8-TBG-*Abhd6* as controls. One-week post-injection, mice were challenged with a 12-week HFD. LABHD6 S148A OE and LABHD6 OE resulted in comparable impairments in pyruvate tolerance relative to control mice, without affecting body weight (Fig. 3M-N). Based on these results, we concluded that the effects of ABHD6 on insulin resistance are unlikely to be mediated either by ABHD6 enzyme activity or by changes in specific lipid species.

### ABHD6 is a pivotal regulator of FoxO signaling

To identify the potential mechanism(s) mediating the effects of ABHD6 on MASLD development, we performed bulk RNA sequencing of livers collected from LABHD6 OE and littermate control mice. Hepatocyte ABHD6 overexpression resulted in a significant upregulation of 131 genes, including *Pck1* and *G6pc*, and downregulation of 165 genes (Fig. S5A). Pathway analysis indicated many pathways, including steroid biosynthesis, the FoxO signaling, lysosome, and others, have been significantly changed (Sup Fig. 5B-C). Of all the pathway, we were participially interested in FoxO signaling, as it was highly correlated with our observed metabolic phenotypes (Sup Fig. 5B-E). We furthermore analyzed genes associated with FoxO signaling and found that most of these genes were affected by ABHD6 overexpression (Fig. S5F). These results strongly support the idea that ABHD6 exerts its effects by regulating FoxO signaling.

**Figure 4:**
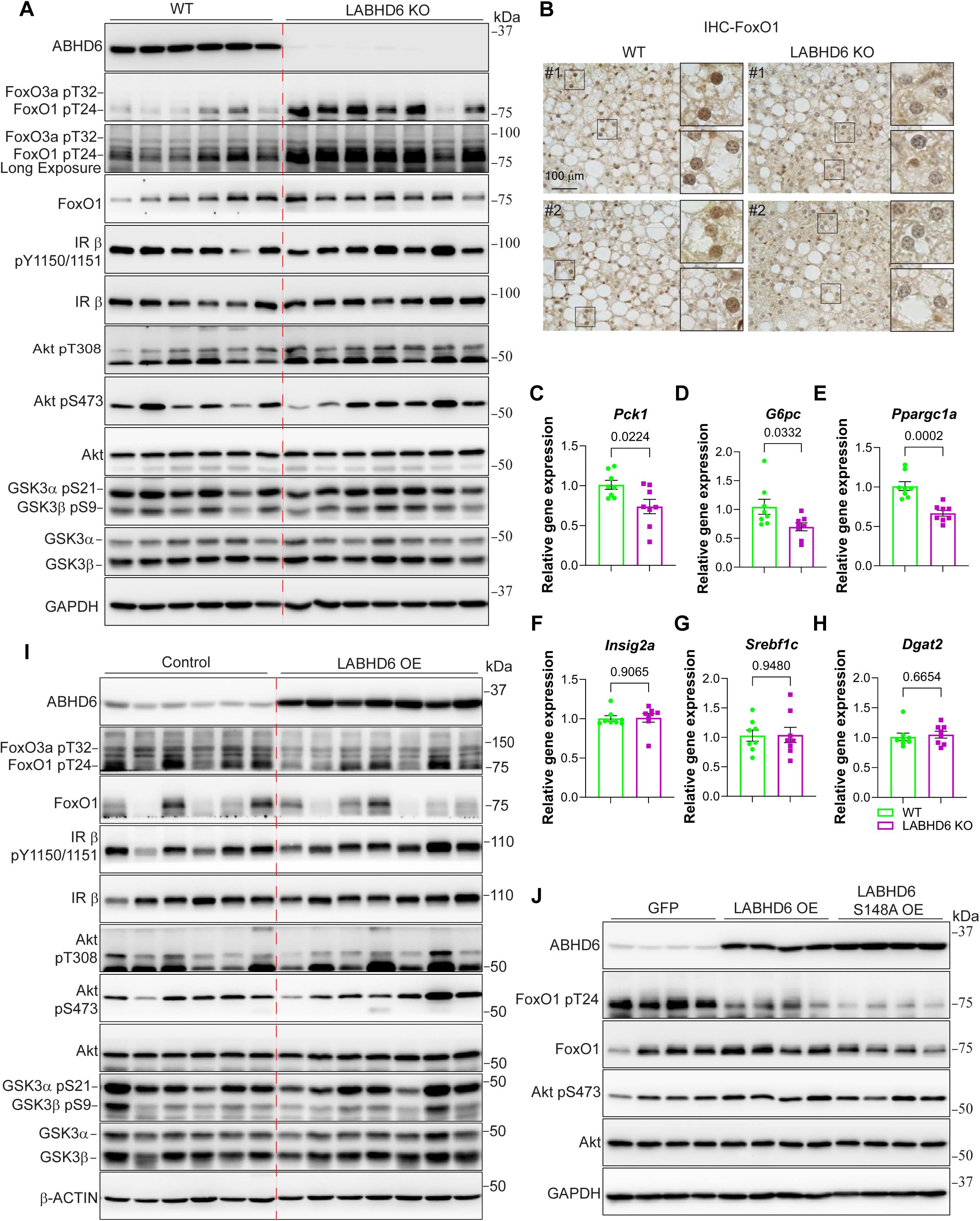
ABHD6 manipulation drives selective hepatic insulin resistance. **(A-H)** 8-week-old littermate control (WT) and LABHD6 KO mice were placed on high-fat diet (HFD) for 8 weeks. **(A)** Protein levels of insulin signaling components in the liver (n = 6-7 per group). **(B)** IHC staining of FoxO1 in the livers of WT and LABHD6 KO mice (n = 2 per group). **(C-H)** mRNA levels of gluconeogenesis and lipogenesis genes in the livers (n = 8 per group). **(I-J)** 8-week-old male wildtype mice were injected with 1×10¹¹ genome copies AAV 8 TBG-eGFP (Control), TBG-*Abhd6* (LABHD6 OE) or TBG-*Abhd6* p.S148A (LABHD6 S148A OE). One-week post-injection, the mice were fed a high-fat diet (HFD) for 12 weeks. **(I)** Protein levels of insulin signaling components in the livers (n = 6-7 per group). **(J)** Protein levels of insulin signaling components in the livers (n = 4 per group). (C-H) Data are displayed as mean ± SEM and analyzed by Student’s *t* test. p-values of respective comparisons are provided.

**Figure 5:**
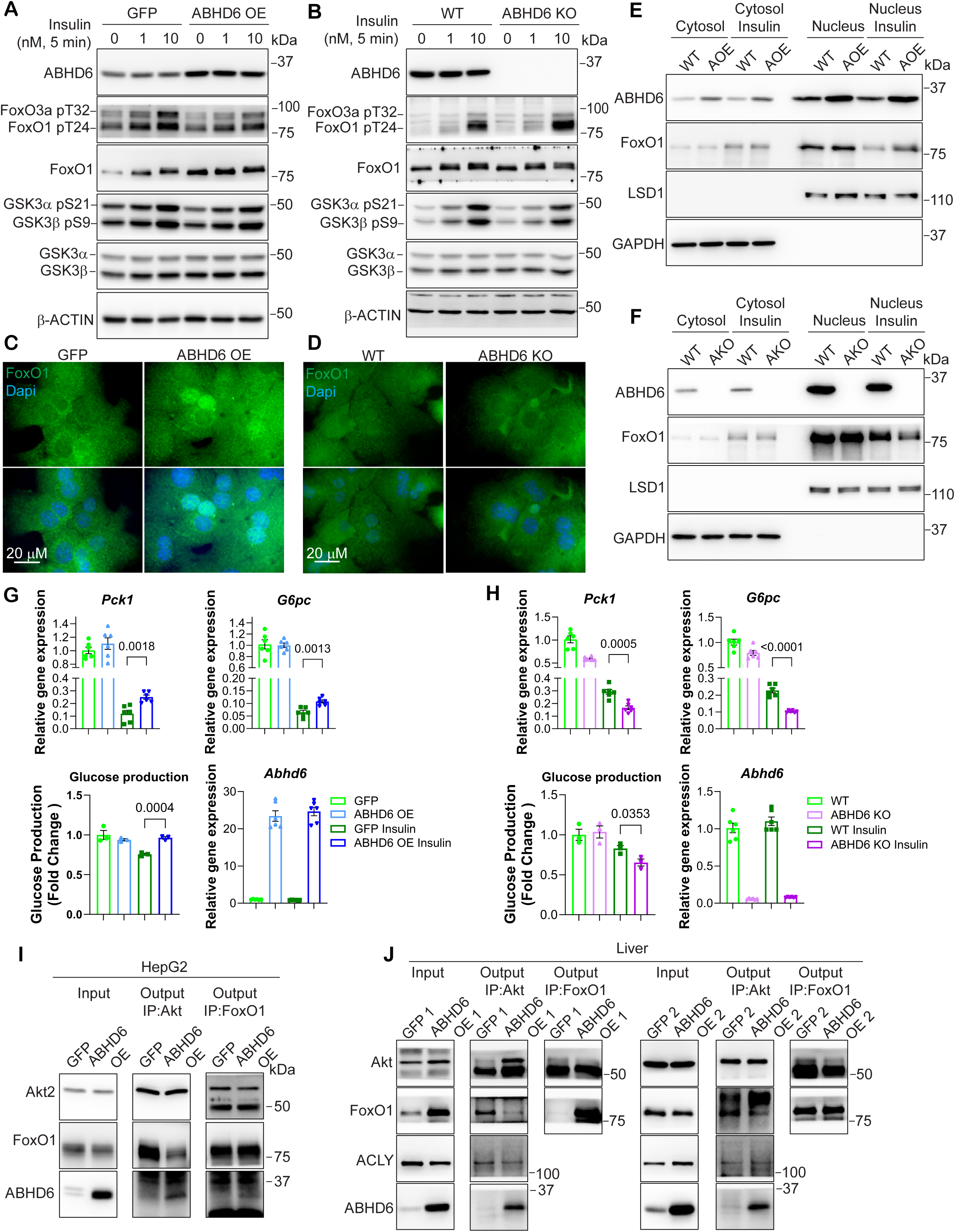
ABHD6 negatively regulates the interaction of Akt with FoxO1. **(A, C, E, G)** Primary hepatocytes were isolated from 8-12-week-old male wildtype mice and transduced with AAVDJ CAG-*Abhd6* (ABHD6 OE) or CAG-eGFP (GFP) at a multiplicity of infection (MOI) of 100. Analyses were performed 48 hours post-transduction. (**B, D, F, H**) Primary hepatocytes were isolated from 8-12-week-old male control littermate (WT) and LABHD6 KO (ABHD6 KO) mice. **(A and B)** Protein levels of ABHD6 and insulin signaling components in primary hepatocytes following insulin stimulation. **(C and D)** FoxO1 immunofluorescence staining showing protein translocation in primary hepatocytes following insulin stimulation. **(E and F)** Nucleus fractionation showing Foxo1 translocation primary hepatocytes following insulin stimulation. AOE, ABHD6 OE; AKO, ABHD6 KO. **(G and H)** mRNA levels gluconeogenic genes, glucose production, and mRNA levels of *Abhd6* in primary hepatocytes. **(I)** HepG2 cells were transfected with 1 mg HA-Akt2 and FLAG-FoxO1 plasmids and transduced with AAVDJ CAG-*Abhd6* (ABHD6 OE) or CAG-eGFP (GFP) at MOI of 1000. 48 hours post-transfection/transduction, Akt2 and FoxO1 co-immunoprecipitation was performed. Protein levels of Akt2, FoxO1, and ABHD6 in input and output samples were determined. **(J)** Akt and FoxO1 co-immunoprecipitations were performed using liver samples from Fig. 3. Protein levels of Akt2, FoxO1, ACLY, and ABHD6 in input and output samples were determined (n = 2 per group). (G and H) Data are displayed as mean ± SEM and analyzed by Student’s *t* test. p-values of respective comparisons are provided.

Expanding on this, we also performed bulk RNA sequencing of mouse primary hepatocytes that we transfected with siRNA targeting *Abhd6*. In this case, we selected primary hepatocytes instead of using liver tissues to minimize transcriptional noise arising from non-parenchymal cells such as Kupffer cells and hepatic stellate cells. Prior to RNA-seq, the success of the siRNA-mediated *Abhd6* knockdown in the cells was confirmed (Sup Fig. 5G). Our bulk RNA sequencing yielded 126 upregulated and 129 downregulated genes (Fig. S5H) and pathway analyses indicated several distinct pathways to be altered (Fig. S5I). Comparing the top 20 pathways each, we found ABHD6 overexpression (Sup Fig. 5A-F) and knockdown (Sup Fig. 5G-I) both affect FoxO signaling (Fig. S5C and S5I), suggesting FoxO signaling to be a crucial mediator of ABHD6’s effects on MASLD progression.

### ABHD6 manipulation drives selective hepatic insulin resistance

As bulk RNA sequencing suggests that ABHD6 regulates FoxO signaling, we next asked whether this regulation involves changes in FoxO protein post-translational modification and/or protein stability. To this end, we analyzed liver samples from both LABHD6 KO and LABHD6 OE mice.

Immunoblot analyses revealed a drastic increase in FoxO1 (Thr24) phosphorylation and a moderate increase in FoxO3a (Thr32) phosphorylation in the livers of LABHD6 KO mice (Fig. 4A). The increase in FoxO1 (T24) phosphorylation resulted in decreased nuclear localization of FoxO1 as previously reported^34^ (Fig. 4B). Consequently, we also detected a significant reduction in the mRNA expression of gluconeogenic genes downstream of FoxO1/3a in the liver of LABHD6 KO mice, including *Pck1*, *G6pc*, and *Ppargc1a* (*Pgc1*α) (Fig. 4C-E). Besides FoxO1/3a, we examined other key targets in insulin signaling. To our surprise, we found that the phosphorylation of the insulin receptor (IRβ Tyr1150/1151), Akt (Thr308 and Ser473), and the Akt substrate GSK3α/β (Ser21/9) were comparable between LABHD6 KO and littermate control livers (Fig. 4A). Additionally, the phosphorylation of key lipogenic regulators downstream of Akt such as ACLY (Ser455) and activation of mTORC1 (indicated by PRAS40 (Thr246), mTOR (Ser2448), and 4E-BP1 (Thr37/46) phosphorylation) was largely unchanged in the livers of LABHD6 KO mice (Fig. S6A). Furthermore, the mRNA expression of *Insig2a*, which usually is suppressed by Akt to promote SREBP1 processing and lipogenesis^15^, was not affected by the elimination of ABHD6 (Fig. 4F). Similarly, the mRNA expression of other lipogenic genes remained unchanged, including *Srebf1*, *Dgat2*, *Acly*, *Acss2*, *Acaca1*, *Fasn*, *Scd1*, and *Insig1*, except for a significant decrease in *Dgat1* (Fig. 4G-H and S6B).

Immunoblot analyses of LABHD6 OE and littermate control livers revealed reduced FoxO1 (Thr24) and FoxO3a (Thr32) phosphorylation upon ABHD6 overexpression, while the phosphorylation of the insulin receptor (IRβ Tyr1150/1151) and Akt (Thr308 and Ser473) remained largely unchanged (Fig. 4I). Additionally, the phosphorylation of key Akt substrates such as GSK3α/β (Ser21/9), ACLY (Ser455), PRAS40 (Thr246) and mTORC1 activation (indicated by mTOR (Ser2448), and 4E-BP1 (Thr37/46) phosphorylation) was largely unaffected in LABHD6 OE mice (Fig. 4I and S6C). In line with these changes in protein phosphorylation, the mRNA expression of the FoxO1/3a-regulated gluconeogenic genes *Pck1* and *G6pc* was increased in the liver of LABHD6 OE mice (Fig. S6D), and that of *Ppargc1a* (*Pgc1*α) showed trend toward increase (Fig. S6D). At the same time, the mRNA expression of lipogenic genes remained unchanged (Fig. S6E). Taken together, these results demonstrate that ABHD6 selectively modulates the activity of FoxO1 and FoxO3a in hepatocytes. Our results suggest that elevated ABHD6 levels selectively diminish the suppressive effect of insulin signaling on gluconeogenesis without affecting its stimulating effect on lipogenesis, thus directly contributing to the development of selective hepatic insulin resistance in MASLD.

### ABHD6 negatively regulates the interaction of Akt with FoxO1

To gain further insights into how ABHD6 regulates the phosphorylation of FoxO1 and FoxO3a, we performed a comprehensive set of *in vitro* experiments using mouse primary hepatocytes with either ABHD6 overexpression (ABHD6 OE) or ABHD6 knockout (ABHD6 KO). In ABHD6 OE primary hepatocytes, the insulin-induced phosphorylation of FoxO1 (Thr24) and FoxO3a (Thr32) by Akt was largely diminished but not that of GSK3α/β (Ser21/9) (Fig. 5A). In line with our previous observations, the phosphorylation of the insulin receptor (IRβ Tyr1150/1151), Akt (Thr308 and Ser473), ACLY (Ser455), and PRAS40 (Thr246) were not affected by ABHD6 overexpression (Fig. S7A). In ABHD6 KO primary hepatocytes, in turn, the insulin-induced phosphorylation of FoxO1 (Thr24) and FoxO3a (Thr32) was elevated (Fig. 5B), whereas that of GSK3α/β (Ser21/9), the insulin receptor (IRβ Tyr1150/1151), Akt (Thr308 and Ser473), ACLY (Ser455), and PRAS40 (Thr246) was unaffected (Fig. 5B and S7B).

Immunofluorescence staining indicated that the decreased FoxO1 phosphorylation resulted in FoxO1 retention in the nucleus of ABHD6 OE primary hepatocytes, while in ABHD6 KO primary hepatocytes, the increased FoxO1 phosphorylation promoted its exclusion from the nucleus in response to insulin treatment (Fig. 5C-D). Nuclear fractionation confirmed the negative role of ABHD6 on insulin-induced FoxO1 nucleus export without affecting nuclear Akt activity and further revealed that ABHD6 co-localized with Akt and FoxO1 in the nucleus (Fig. 5E-F and Fig.S7D). Consequently, ABHD6 OE impaired the insulin-mediated suppression of gluconeogenic gene expression, including that of *Pck1* and *G6pc*, as well as glucose production in primary hepatocytes (Fig. 5G). Complementing these observations, ABHD6 KO enhanced the insulin-mediated suppression of gluconeogenic gene expression and glucose production in primary hepatocytes (Fig. 5H).

To investigate how ABHD6 impacts the interaction of Akt with FoxO1, we performed immunoprecipitation assays in HepG2 cells overexpressing ABHD6 (ABHD6 OE). In these cells, ABHD6 was pulled down by both Akt2 and FoxO1, and ABHD6 OE substantially weakened the Akt2-FoxO1 interaction (Fig. 5I). ABHD6 OE did not affect FoxO1-PP2A interaction^36^, suggesting ABHD6 likely regulates FoxO1 phosphorylation by Akt rather than its dephosphorylation (Fig. S7E). Moreover, the presence of both Akt2 and FoxO1 strengthened the interaction between Akt2 and ABHD6 (Fig. S7F) compared to Akt2 alone, suggesting ABHD6 may participate in a ternary interaction with Akt2 and FoxO1. We furthermore assessed the endogenous Akt-FoxO1 interaction in the livers of LABDH6 OE mice. Doing so, we found that the Akt-FoxO1 interaction was suppressed by ABHD6 overexpression, using either FoxO1 or Akt as bait, whereas the interaction between Akt and ACLY, another Akt substrate^37^, remained largely unchanged (Fig. 5J). Thus, ABHD6 colocalizes with Akt and FoxO1 in the nucleus and negatively and selectively regulates the Akt-FoxO1 interaction, thereby preventing the Akt-mediated phosphorylation and subsequent nuclear export of FoxO1.

### ABHD6 regulates insulin sensitivity in a FoxO1-dependent manner

To verify the negative regulatory role of ABHD6 in FoxO1 phosphorylation and insulin sensitivity, we asked whether inhibition of FoxO1 could rescue the insulin resistant phenotype of LABHD6 OE mice. To this end, we orally administered FoxO1 inhibitor AS1842856 in LABHD6 OE mice as previously described^35^. Inhibition of FoxO1 activity by AS1842856 improved pyruvate and glucose tolerance in LABHD6 OE mice challenged with HFD without affecting bodyweight (Fig. 6A-B). Accordingly, gene expression of *Pck1* was suppressed by FoxO1 inhibition (Fig. S8A). Moreover, an siRNA-mediated knockdown of *Foxo1* largely restored the suppression of gluconeogenic gene expression by insulin in ABHD6 OE primary hepatocytes (Fig. S8B). In addition, we generated AAV8-TBG-*Foxo1* p.T24, which expresses an Akt-phosphorylation-deficient and thus constitutively active FoxO1 mutant (T24A). Overexpression of FoxO1-T24A in LABHD6 KO mice abrogated the insulin-sensitizing effect of ABHD6 KO in pyruvate and glucose tolerance test without affecting bodyweight (Fig. 8D-F). Thus, ABHD6 functions as an upstream regulator of FoxO1 activity, thereby contributing to insulin resistance.

**Figure 6:**
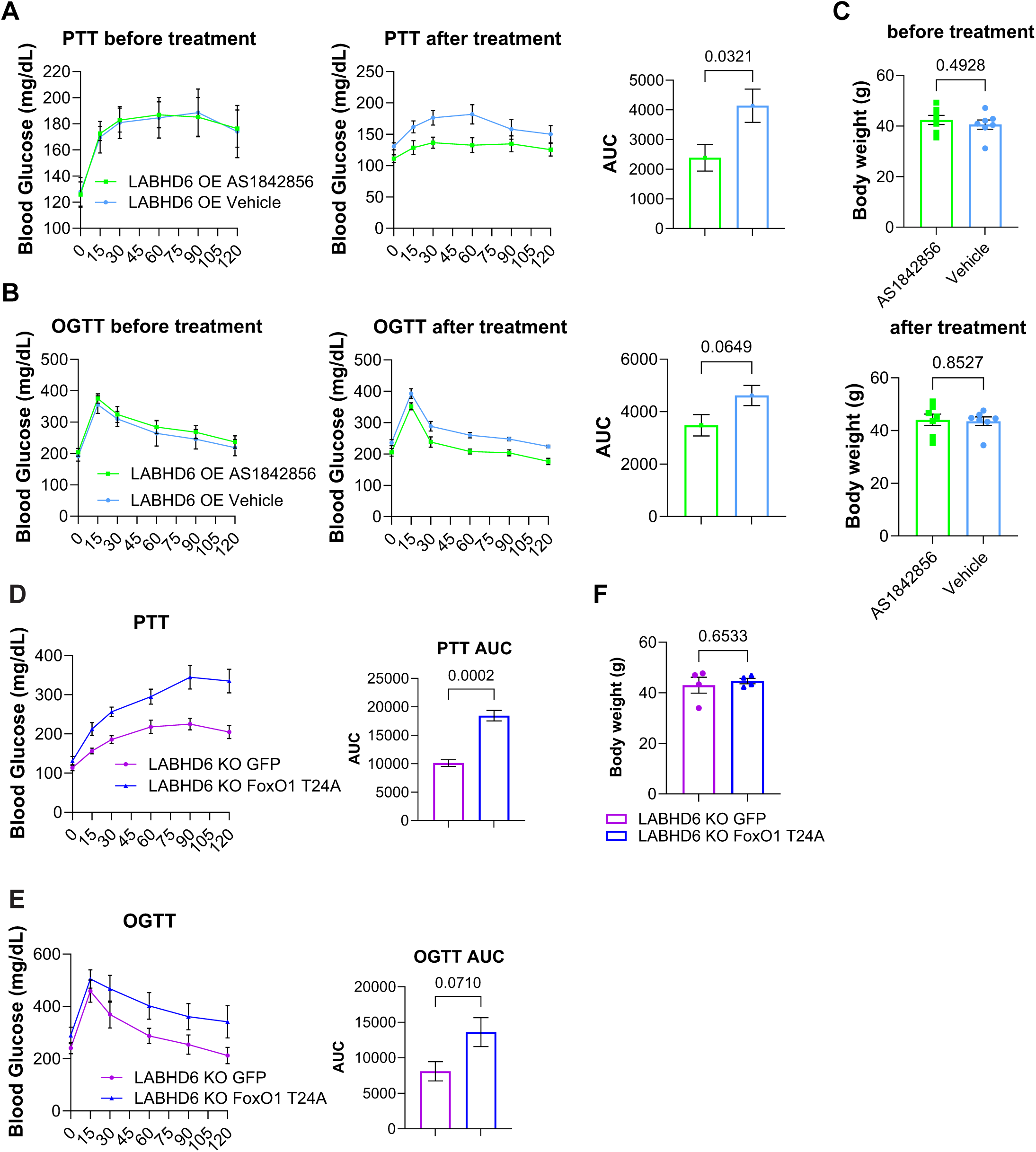
ABHD6 regulates insulin sensitivity in a FoxO1-dependent manner. **(A-C)** 8-week-old male wildtype mice were injected with 1×10¹¹ genome copies TBG-Abhd6 (LABHD6 OE). One-week post-injection, the mice were fed a high-fat diet (HFD) for 12 weeks AS1842856 was administered orally at 30 mg/kg, three times over two days prior to PTT and GTT (see Methods for experimental details). **(A and B)** Blood glucose levels during pyruvate tolerance test (PTT) and glucose tolerance test (GTT). For the PTT and GTT, the area under the curve (AUC) was calculated (n = 7 per group). **(C)** Body weight before and after AS1842856 treatment (n = 7 per group). **(D and E)** Blood glucose levels during pyruvate tolerance test (PTT) and glucose tolerance test (GTT). For the PTT and GTT, the area under the curve (AUC) was calculated (n=4 per group). **(F)** Body weight (n = 4 per group).

## Discussion

In this study, we identify ABHD6 as a novel contributor to MASLD establishment and progression. We demonstrate that ABHD6 promotes selective hepatic insulin resistance by interfering with the interaction of Akt with FoxO1/3a. Moreover, we demonstrate that this effect of ABHD6 on liver function is mainly contributed by its non-enzymatic function. In addition, we found the effects of ABHD6 is completely dependent on its interactions with Akt-FoxO1 axis.

The enzymatic function of ABHD6 has been well documented. Previously, we demonstrated that ABHD6-accessible monoacylglycerol is a new metabolic coupling factor that regulates glucose-stimulated insulin secretion by binding to Munc13-1^31,38^. In addition, using antisense oligonucleotide (ASO) to knockdown ABHD6 in multiple tissues, including liver, white adipose tissue, brown adipose tissue, kidney and spleen, Dr. Brown group demonstrated that ABHD6 is a critical factor in the establishment of insulin resistance by regulating *de novo* lipogenesis^39^. Furthermore, by generating global ABHD6 deficient mice, as well as adipocytes-specific ABHD6 knockout mice, we found that inhibition of ABHD6 in adipose tissue induces browning of white adipocytes and enhances their thermogenic activity^32,33^. The effects of ABHD6 on adipose tissue explains why global ABHD6 KO and ASO-mediated ABHD6 knockdown mice are protected from diet-induced obesity, leading to a much-improved glucose tolerance and insulin sensitivity. Recent studies also suggest an important role of ABHD6 in regulating food intake in the brain *via* 2-AG^40^. All these studies indicate the importance of the enzymatic function of ABHD6. However, none of these studies explored the non-enzymatic function of ABHD6. Furthermore, the observed beneficial effects on glucose tolerance and insulin sensitivity in global ABHD6 KO and ASO-mediated ABHD6 knockdown are thought to be mediated by knockdown of ABHD6 in adipose tissue. Previously, we demonstrated that deletion of ABHD6 in adipose tissue induces browning of white adipose tissue and enhances brown fat function, leading to weight-loss and protection from MASLD. Therefore, the present study represents the first systematic investigation of ABHD6 function specifically in the liver, as hepatocyte-restricted ABHD6 deletion and overexpression produce phenotypes that are distinct from those observed in global or ASO-mediated ABHD6 loss. Interestingly, hepatocyte-specific ABHD6 overexpression reduced hepatic triglyceride accumulation despite markedly exacerbating insulin resistance, inflammation, and fibrosis. While the reduction in hepatic lipid content is consistent with the canonical enzymatic function of ABHD6 in lipolysis, the severe metabolic and fibrotic phenotypes cannot be explained by this activity alone, highlighting a dominant non-enzymatic role for ABHD6 in disease progression.

Based on the comprehensive lipidomic analysis in our ABHD6 manipulated mice, together with phenotypic characterization of mice expressing an enzymatically inactive ABHD6 mutant, we identified a non-enzymatic function of ABHD6 that contributes to the major metabolic phenotypes in this study, including glucose tolerance, insulin sensitivity and gluconeogenesis. Using nuclear fractionation and immunofluorescence analyses, we further observed that ABHD6 is translocated into the nucleus, where immunoprecipitation studies revealed its association with the Akt–FoxO1 signaling axis, resulting in enhanced FoxO1 transcriptional activity. The identification of a nuclear, non-enzymatic role for ABHD6 in transcriptional regulation reveals a previously unrecognized mechanism by which ABHD6 influences metabolic homeostasis and opens new avenues for investigating its role in metabolic disease. Whether nuclear ABHD6 also regulates the activity of additional transcription factors remains an important question for future studies. Previous reports have indicated that ABHD6 promotes the endocytosis of AMPA receptors to regulate synaptic plasticity and learning flexibility^41,42^. Similarly, our findings suggest that nuclear ABHD6 may exert regulatory functions in transcriptional control beyond its canonical catalytic role.

Great efforts have been made to elucidate the potential mechanisms underlying selective hepatic insulin resistance. Here, we propose obesity-associated increases in ABHD6 expression and the resulting selective suppression of FoxO signaling in hepatocytes to play a crucial role. Through this action of ABHD6, gluconeogenesis in particular is spared from insulin signaling-dependent suppression. We found that ABHD6 is increased in liver during diet-induced obesity and aging. In addition, we detected ABHD6 in nuclear extracts, which may support the retention of FoxO1/3a in the nucleus to actively transcribe gluconeogenesis genes such as *Pck1* and *G6pc*. This idea is strongly supported by our experimental data. As such, the overexpression of ABHD6 in hepatocytes causes a disruption of the Akt-FoxO1 interaction and blocks the trafficking of FoxO1 from the nucleus to the cytoplasm. In contrast, the elimination of ABHD6 from hepatocytes promotes the nucleus-to-cytoplasm trafficking of FoxO1, thus effectively diminishing the transcriptional activity of FoxO1 and gluconeogenesis gene expression. In addition, knockdown of FoxO1 or overexpression of mutant FoxO1 completely abolished ABHD6’s effect on glucose tolerance and gluconeogenesis, indicating the importance of the interaction between ABHD6 and the Akt-Foxo1 axis. These effects of ABHD6 on FoxO signaling provide a new angle to explain the establishment of selective hepatic insulin resistance.

Overexpression of ABHD6 in hepatocytes caused a severe deterioration of systemic metabolism, characterized by pronounced glucose intolerance and insulin resistance, increased liver gluconeogenesis, and increased liver inflammation and fibrosis. However, overexpression of ABHD6 in hepatocytes was not accompanied by an increase in the accumulation of intracellular lipid droplets. This manifestation of liver disease resembles that found in whole-body or hepatic stellate cell-specific adiponectin knockout mice^43,44^. In these mouse models, we previously observed a similar decrease in liver lipid content but concomitant increase liver inflammation and fibrosis^43,44^. In addition, this manifestation of liver disease is commonly seen in cases of chronic hepatitis B or C infection, autoimmune dysfunction, or in certain genetic conditions. Our hepatocyte-specific ABHD6 overexpression mouse model may thus provide a valuable tool for the further study of this specific form of liver disease.

FoxO signaling is implicated in multiple diseases, particularly metabolic disorders like obesity, diabetes, atherosclerosis, and MASLD, as well as B-cell malignancies and potentially neurodegenerative diseases^45–47^. Our finding that non-enzymatic ABHD6 is a regulator of FoxO activity could explain the beneficial effects of ABHD6 inhibition in the settings of obesity, diabetes, MASLD, and liver fibrosis. In addition, the regulation of FoxO signaling by ABHD6 further implies a useful application of ABHD6 inhibition in other settings such as neurodegenerative diseases.

## Material and Methods

### Mouse models

All animal experimental protocols have been approved by the Institutional Animal Care and Use Committee (IACUC) of the University of Texas Health Science Center at San Antonio. Male C57BL/6J mice (4–5 mice per cage) were housed under standard laboratory conditions (12 hours on/off; lights on at 7:00 a.m.) in a temperature-controlled environment with *ad libitum* access to food and water. All experiments were initiated at approximately 8 weeks of age. For all *in vivo* experiments, littermate control mice were used. The mouse genotype had no apparent effect on initial weight, overall health, or immune status of the animals.

To explore the impact of obesity on ABHD6 expression, male mice were fed a high-fat diet (HFD; Bio-Serv, S1850, 60% kcals from fat) or chow diet (control) for 8 weeks. To explore the impact of aging on ABHD6 expression, 18-month-old mice were considered ‘old’ and 6-month-old male mice ‘young’.

Liver-specific ABHD6 knockout (LABHD6 KO) was achieved by crossing *Abhd6* flox mice (Generated in Dr. Marc Prentki’s lab) with Albumin-Cre mice (RRID: IMSR_Jax:003547). The resulting LABHD6 KO mice were then fed HFD, starting at 8-week-old. Liver-specific ABHD6 overexpression (LABDH6 OE) was achieved by tail-vein injection of adeno-associated viruses (AAVs), specifically AAV 8 TBG-*Abhd6* and TBG-eGFP (control), into wildtype mice at a dose of 1×10¹¹ genome copies (gc) per mouse. One week after the injection, the mice were placed on HFD.

After 8 weeks (LABHD6 KO) or 12 weeks (LABHD6 OE) of HFD feeding, oral glucose tolerance tests (GTTs), intraperitoneal insulin tolerance tests (ITTs), and intraperitoneal pyruvate tolerance tests (PTTs) were performed as previous described^48^. Afterwards, all mice were euthanized, and blood was collected from inferior vena cava. Subsequently, the liver was perfused with PBS and liver and adipose tissue weights were determined. Blood samples were allowed to clot at room temperature and centrifuged for 10 min at 2000 g, 4°C. The serum supernatant was collected, flash-frozen in liquid nitrogen and stored at -80°C. Tissue samples were either snap-frozen in liquid nitrogen or fixed in formalin for further analysis.

To inhibit FoxO1 activity in LABHD6 OE mice, AS1842856 was orally administered as previously described^35^. Briefly, 30 mg/kg AS1842856 was administered three times, 9:00 am and 6:00 pm on day 1 and 8:00 am on day 2. For PTT of AS1842856 treated mice, mice were fasted overnight on day 1 after second AS1842856 administration, and PTT was performed 2 hours after last AS1842856 administration. For OGTT, mice were fasted after the last AS1842856 administration and for 6 hours.

### Primary hepatocyte isolation and culture

Mouse primary hepatocytes were isolated from 8- to 12-week-old wildtype or LABHD6 KO mice on a pure C57BL/6J background by retrograde perfusion with liberase described previously^49^. A total of 5x10^5^ cells were seeded in each well of a 6-well-plate. Primary hepatocytes were cultured with DMEM (Corning, 1g/L glucose, 10-014-CV) supplemented with 5% FBS (Corning, 35-011-CV), 100 units/mL of penicillin and 100 μg/mL of streptomycin for 4 hours to allow the cells to attach. Following attachment, the cells were washed with pre-warmed PBS twice and the medium was switched to William’s E medium (Gibco, no phenol red, no FBS, A1217601) supplemented with 2 mM L-glutamine.

In cases where no further treatment was performed, the cells were continuously cultured in William’s E medium overnight, and analyses were conducted the next day. For AAV treatment, the cells were transduced with either AAV DJ CAG-*Abhd6* or CAG-eGFP (control) at a multiplicity of infection (MOI) of 100, concurrent with the change to William’s E medium. Analyses were conducted 48 hours post-transduction. For siRNA treatment, the cells were transfected with siRNA at a final concentration of 20 nM using Lipofectamine 3000 (Invitrogen, L3000015), concurrent with the change to William’s E medium. Analyses were conducted 48 hours post-transfection.

### HepG2 culture

HepG2 cells were grown in DMEM (Corning, 4.5g/L glucose, 10-013-CM) supplemented with 10% FBS (Corning, 35-011-CV) at 37°C, 5% CO_2_ in a humified atmosphere. For concurrent plasmid transfection and AAV treatment, 1x10^6^ cells were seeded in each well of a 6-well-plate. After 24 h, the cells were transfected with 1 µg plasmid encoding HA-Akt2 and FLAG-FoxO1 using Lipofectamine 3000 (Invitrogen, L3000015). AAV DJ CAG-*Abhd6* or CAG-eGFP (control) was added at a MOI of 1000 right after transfection. Analyses were performed 48 hours post-transfection/transduction.

### Protein isolation and immunoblot

Proteins were isolated using RIPA buffer (20 mM Tris-HCl, 1 mM EDTA, 0.5 mM EGTA, 1% Triton X-100, 0.1% sodium deoxycholate, 0.1% sodium dodecyl sulfate, 150 mM NaCl; final pH 7.4) supplemented with protease and phosphatase inhibitors (ThermoFisher Scientific). Tissue samples were homogenized in buffer using a TissueLyer II (Qiagen). For cultured cells, the buffer was directly added to the culture plates, and the cells were collected using a cell scraper. The crude protein extracts were briefly sonicated and incubated for 20 minutes on ice, followed by centrifugation for 10 minutes at 13,000 rpm, 4°C. Following centrifugation, the supernatant was collected, and protein concentrations were measured using the BCA protein assay kit (ThermoFisher Scientific, 23225).

Subsequently, 10-15 µg of total protein was separated by SDS-PAGE and transferred onto nitrocellulose membranes. The following primary antibodies were used: ABHD6 (CST, #97573, 1:1000), FoxO1 (CST, #2880, 1:1000), phospho-FoxO1 (Thr24)/FoxO3a (Thr32) (CST, #9464, 1:1000), phospho-IGF1Rβ (Tyr1135/1136)/INSRβ (Tyr1150/1151) (CST, #3024, 1:1000), INSRβ (CST, #3025, 1:1000), phospho-Akt (Ser473), (CST, #4060, 1:1000), phospho-Akt (Thr308) (CST, #13038, 1:1000), Akt (pan) (CST, #4691, 1:1000), phospho-GSK3α/β (Ser21/9) (CST, #8566, 1:1000), GSK3α/β (CST, #5676, 1:1000), phospho-ACLY (Ser455) (CST, #4331, 1:1000), ACLY (CST, #4332, 1:1000), phospho-PRAS40 (Thr246) (CST, #13175, 1:1000), PRAS40 (CST, #2691, 1:1000), phospho-mTOR (Ser2448) (CST, #5536, 1:1000), mTOR (CST, #2972, 1:1000), phospho-4E-BP1 (Thr37/46) (CST, #2855, 1:1000), 4E-BP1 (CST, #9644, 1:1000), LSD1 (CST, #2184, 1:1000), β-Actin (Abclonal, AC004, 1:10,000), and GAPDH (Abclonal, AC002, 1:10,000).

### Immunofluorescence staining

Primary hepatocytes were cultured on coverslips and treated with 10 nM insulin for 5 minutes before fixation with 4% formaldehyde (FA) for 10 minutes at room temperature. The coverslips were then rinsed three times with PBS and permeabilized with 0.3% (v/v) Triton X-100 in PBS for 10 minutes at room temperature. After an additional three PBS washes, the cells were blocked with 10% goat serum in PBST (0.1% Tween-20) and then incubated overnight at 4°C with an anti-FoxO1 antibody (1:100 in 10% goat serum). The following day, the coverslips were rinsed three times with PBS and incubated for 1 hour at room temperature in the dark with a goat anti-Rabbit IgG (H+L) cross-adsorbed secondary antibody conjugated to Alexa Fluor 488 (1:500). After three more PBS washes in the dark, the coverslips were mounted on slides using ProLong Diamond antifade mountant with DAPI. The slides were cured overnight at 4°C and images were taken on a BZ-X7800E microscope (KEYENCE).

### Histology

Hematoxylin& Eosin (H&E) staining was performed on liver tissue sections as described previously^50^. Liver tissue was rapidly harvested and fixed in phosphate-buffered 10% formalin (Fisher Chemical, Cat#SF100-20) for 2 weeks at room temperature. Fixed liver tissues were then embedded in paraffin, cut into 4 µm sections, and mounted on glass slides. Paraffin-fixed liver sections were subjected to H&E staining and images were taken on a BZ-X800E microscope (KEYENCE).

### RNA isolation, RT-qPCR and RNAseq

Primary hepatocytes were treated with 100 μM cAMP, with or without 10 nM insulin, for 3 hours before harvesting. RNA of tissue and cells was extracted using TRIzol (Invitrogen, Cat #15596026 and purified by isopropanol precipitation. RNA concentrations were determined on a Nanodrop (Thermo Scientific).

For RT-qPCR, a total of 1 µg RNA was reverse transcribed using the Hiscript III 1^st^ Strand cDNA Synthesis kit (gDNA wiper) (Vazyme, R312). mRNA expression levels were quantified by a CFX384 Real-Time PCR Detection System (Bio-Rad). Quantitative analysis was performed using the ΔΔCt method. The relative mRNA expression levels of genes were normalized to those of 16S. Primers used are listed in Table S1. RNA sequencing was performed by Novogene.

### Nuclear fractionation

The nuclear fraction of primary hepatocytes was prepared using the Nuclei Isolation Kit (Millipore/Sigma, NUC-101). Briefly, primary hepatocytes were treated with 10 nM insulin for 10 minutes, then rinsed with ice-cold PBS. To each well of a 6-well plate, 500 µl of ice-cold Nuclei EZ lysis buffer were added, and cells were gently collected using a cell scraper. After collection, the cell suspension was briefly vortexed and incubated for 5 minutes on ice. Nuclei were then pelleted by centrifugation for 5 minutes at 500 g, 4°C, and the pellet was retained for subsequent washes. The supernatant was carefully collected and centrifuged again for 5 minutes at 500 g, 4°C. The supernatant resulting from this step was saved as the cytosolic fraction. The nuclei pellet from the first centrifugation was resuspended in 300 µL of ice-cold Nuclei EZ lysis buffer by vertexing at moderate to high speed. An additional 700 µL of ice-cold Nuclei EZ lysis buffer was then added, the samples mixed thoroughly and incubated for 5 minutes on ice. Nuclei were collected by centrifugation as described previously, and the supernatant was carefully removed. This wash step was repeated once more. Finally, 200 µL of 2X Laemmli SDS buffer (Bio-Rad, CAT #1610737) were added to the nuclei pellet, followed by boiling for 5 minutes at 95°C to obtain the nuclear fraction.

### Immunoprecipitation

HepG2 cells were cultured in 60 mm dishes, transfected with 1 μg HA-Akt2 and FLAG-FoxO1 plasmid and transduced with AAVdj-CAG-GFP or ABHD6 at MOI1000. 48 hours post transfection/transduction, HepG2 cells were serum-starved for 4 hours, followed by treatment with 10 nM insulin for 5 minutes. HepG2 cells and liver samples were lysed in NP-40 lysis buffer (50 mM Tris-HCl, 150 mM NaCl, 0.5% NP-40; final pH 7.4) supplemented with protease and phosphatase inhibitors (ThermoFisher Scientific). ed with protease and phosphatase inhibitors (ThermoFisher Scientific). Tissue samples were homogenized in buffer using a TissueLyer II (Qiagen). For cultured cells, the buffer was directly added to the culture plates, and the cells were collected using a cell scraper. The crude protein extracts were briefly sonicated and incubated for 20 minutes on ice, followed by centrifugation for 10 minutes at 13,000 rpm, 4°C. Following centrifugation, the supernatant was collected, and protein concentrations were measured using the BCA protein assay kit (ThermoFisher Scientific, 23225).

Protein extracts were pre-cleared by incubation with Protein A Dynabeads, rotating for 2 hours at 4°C. After centrifugation for 5 minutes at 500 g, 4°C, the supernatant was transferred to a new tube, and 1/10 of the supernatant was set aside as an input sample. For immunoprecipitation, 200 µl of liver lysate (∼1.5 mg total protein) or 200 µl of HepG2 lysate (∼1 mg total protein) were incubated overnight rotating at 4°C with primary antibodies against Akt (pan) (CST, #4691, 1:50) and FoxO1 (CST, #2880, 1:100).

The following day, 20 µl of Protein A Dynabead slurry was added to the lysate, and the sample was incubated for 1 hour rotating at 4°C. Beads were then collected by centrifugation for 5 minutes at 500 g, 4°C, and the supernatant was saved as the flow-through fraction. The beads were sequentially washed once with low-salt wash buffer (50 mM Tris-HCl, 150 mM NaCl, 0.5% NP-40), twice with high-salt wash buffer (50 mM Tris-HCl, 500 mM NaCl, 0.5% NP-40), and once more with low-salt wash buffer. Finally, 50 µl of 2X Laemmli SDS buffer were added to the beads, followed by boiling for 5 minutes at 95°C to obtain the output sample.

### *in vitro* gluconeogenesis assay

Primary hepatocytes were cultured in 6-well plates in William’s E medium, washed twice with pre-warmed PBS, and then cultured in glucose production medium (DMEM without glucose, L-glutamine, pyruvate, and phenol red; Millipore/Sigma, D5030) supplemented with 10 mM glycerol. The cells were treated with 100 µM cAMP, with or without 10 nM insulin, for 4 hours. Following treatment, the medium was collected and centrifuged for 4 minutes at 500 g, 4°C. Meanwhile, cells were lysed in RIPA lysis buffer and protein concentrations were measured using the BCA protein assay kit. Glucose levels in the cell culture supernatants were determined immediately after collection using the Autokit Glucose (Fujifilm, 997-03001) and normalized to the determined protein concentrations.

## Data Availability

Source data will be provided with this paper prior to publication. The lipidomic dataset and RNAseq data will be uploaded prior to publication. All quantitative data underlying the figures and Supplementary Figures will be provided prior to publication.

## Abbreviations

ABHD6: α/β hydrolase domain-containing 6
HFD: high-fat diet
GTT: glucose tolerance test
ITT: insulin tolerance test
PTT: pyruvate tolerance test.

## Acknowledgements

This work was supported by US National Institutes of Health grants R00-AG068239, R01-DK138035, and R01-AG084646 to S.Z.; R01-DK136619 and R01-DK136532 to Y.Z.; and R01-DK136848 to J.B. It was furthermore supported by a Hevolution Fund grant (23-1199262-27) and a Voelcker Fund Young Investigator grant to S.Z. Functional lipidomics Core at the Barshop Institute was partially supported by NIA P30 AG013319 and P30 AG044271.

## Author contributions

G.L. and S.Z. designed the study. G.L., L.M., Z. X., and S.Z. analyzed the results. G.L. and S.Z. drafted the manuscript. G.L., L.M., J.Y., JB. F., C.J., R.L., Z.X., T.W., X.L., N.J., M.E., P.J., and M.L. contributed to the acquisition and analysis of data. JB. F, S.H., A. S., M.M., M.P., J.B., J.N., X.H., Y.Z., and S.Z. revised and approved the final version of the manuscript. S.Z. is the guarantor of this work, with full access to all data and analyses related to the content of this article.

## Conflict of interest statement

The authors declare no conflict of interest.

## Supplementary information

A. Supplementary Figures.

1. High fat diet induces liver insulin resistance.

2. Deletion of ABHD6 in liver protects the mice from MASLD and liver fibrosis.

3. Overexpression of ABHD6 in the liver drives insulin resistance through a lipid-independent mechanism.

4. ABHD6 acts as a BMP hydrolase yet modulates insulin signaling independently of BMP.

5. ABHD6 is a pivotal regulator of FoxO signaling.

6. ABHD6 manipulation only affects Akt-Foxo1/3a-mediated gluconeogenesis, without affecting Akt-mediated lipogenesis.

7. ABHD6 regulates Akt-Foxo1/3a phosphorylation and interaction, leading to altered gluconeogenesis.

8. ABHD6 regulates insulin sensitivity in a FoxO1-dependent manner.

B. Supplementary Table

1. Primers used in the study.

2. Antibodies used in the study.

3. DEG pathway in response to ABHD6 manipulation.

---

**Sup Figure 1:**
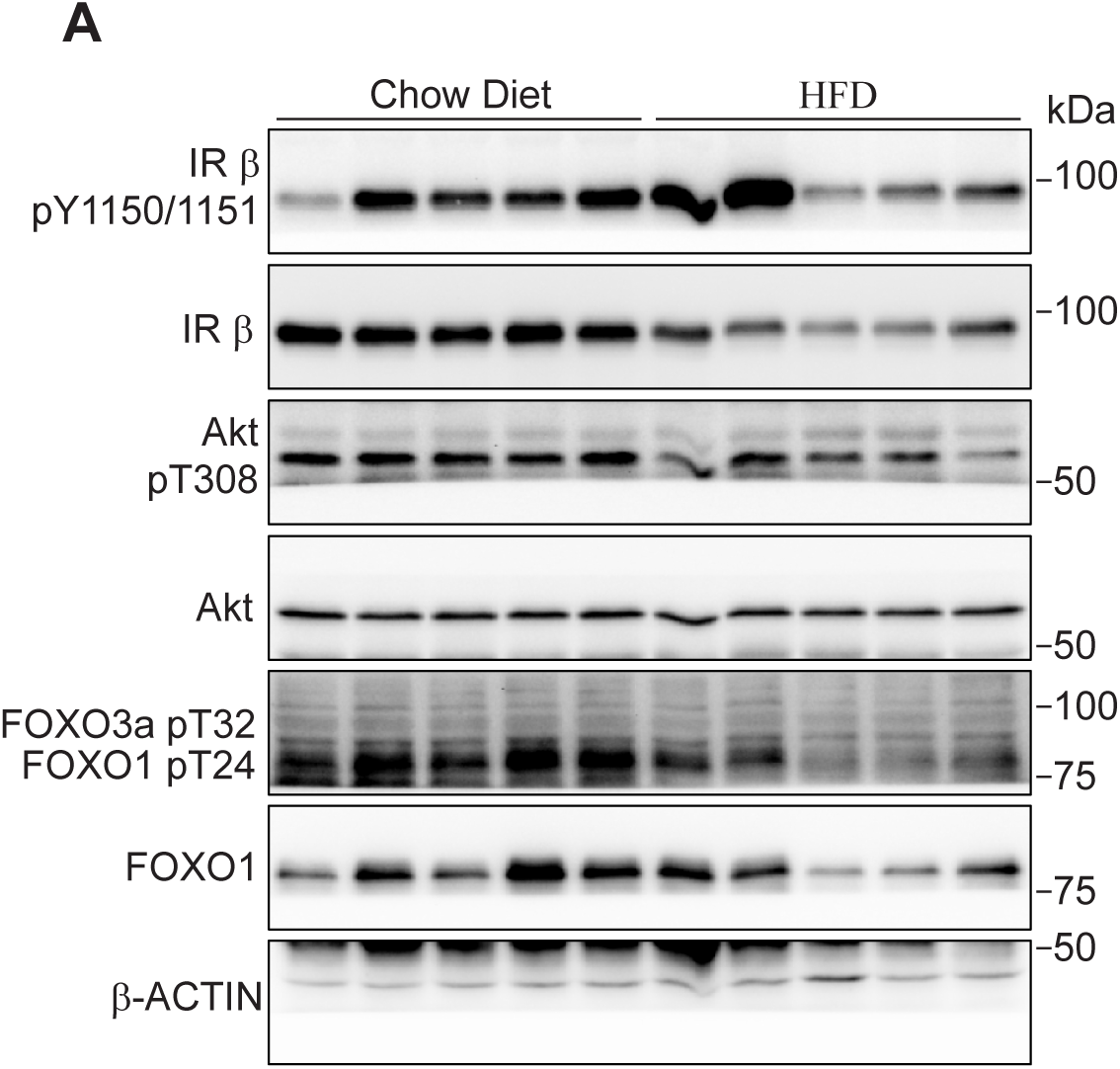
High fat diet induces liver insulin resistance. (A) Effects of high fat diet on hepatic insulin signaling by western blots.

**Sup Figure 2.**
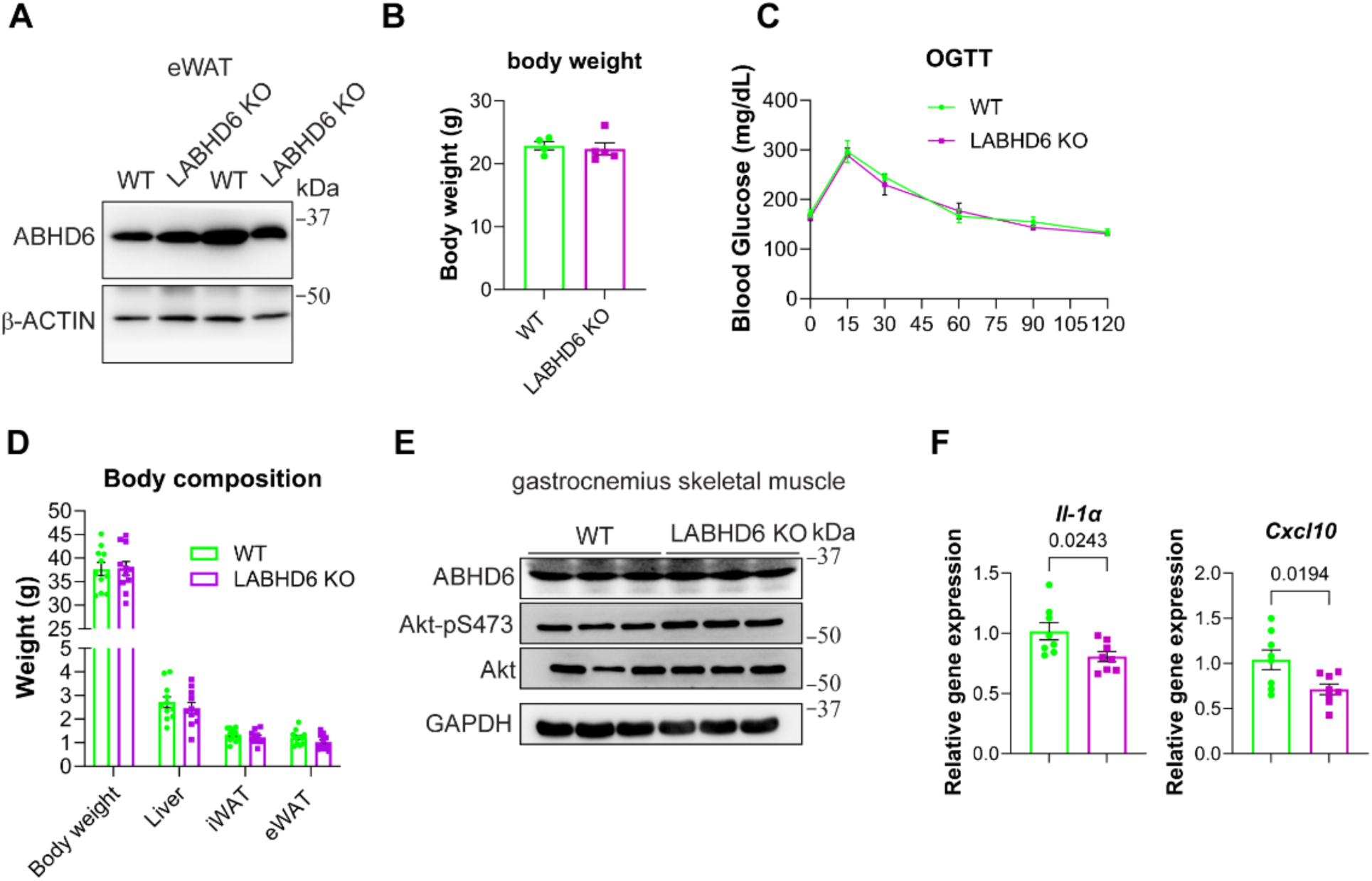
Deletion of ABHD6 in liver protects the mice from MASLD and liver fibrosis. (A) ABHD6 protein levels in adipose tissue. (B) Body weight of WT and LABDH6 KO mice under chow diet. (C) Blood glucose levels during glucose tolerance test (GTT) (n=5 per group). (D) Body composition. (E) Immunoblot of ABHD6 and Akt activation in the gastrocnemius skeletal muscle of LABHD6 KO mice. Mice were deprived of food overnight and intraperitoneal injected with 0.25 U/kg insulin. Gastrocnemius skeletal muscle was harvested 5 minutes after insulin injection. (F) Expression of Il-1a and Cxcl10. Data are displayed as mean ± SEM and analyzed by Student’s *t* test. p-values of respective comparisons are provided

**Sup Figure 3.**
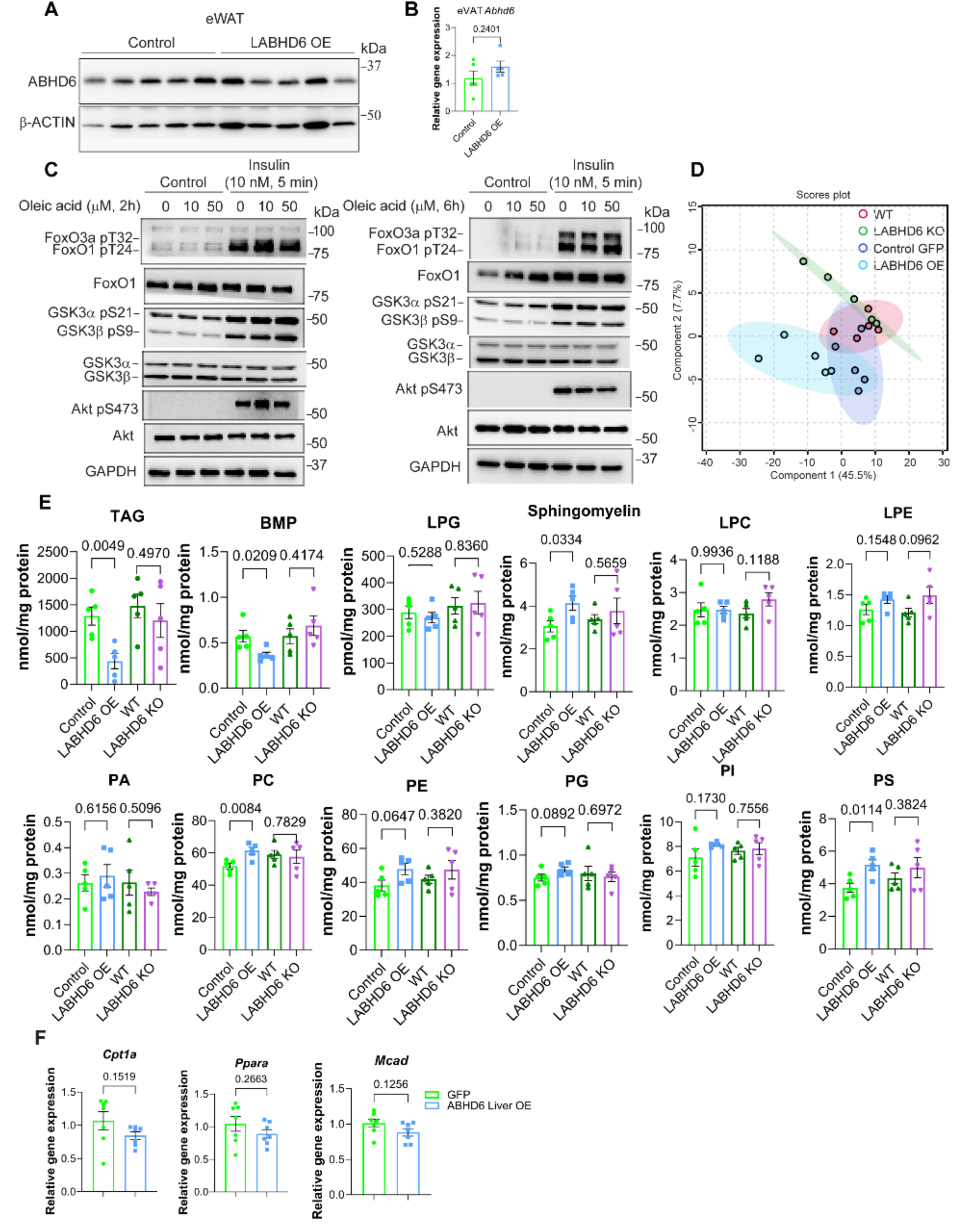
Overexpression of ABHD6 in the liver drives insulin resistance through a lipid-independent mechanism. (A) ABHD6 protein levels in adipose tissue. (B) Gene expression of *Abhd6* in adipose tissue. (C) Effects of exogenous 1-OG administration on hepatic insulin signaling. (D) partial least squares discriminant analysis (PLS-DA). (E) Overview of the change of lipid classes in response to ABHD6 deletion and overexpression. (F) Gene expression of Cpt1a, Ppara, and Mcad. (B, E-F)Data are displayed as mean ± SEM and analyzed by Student’s *t* test. p-values of respective comparisons are provided.

**Sup Figure 4.**
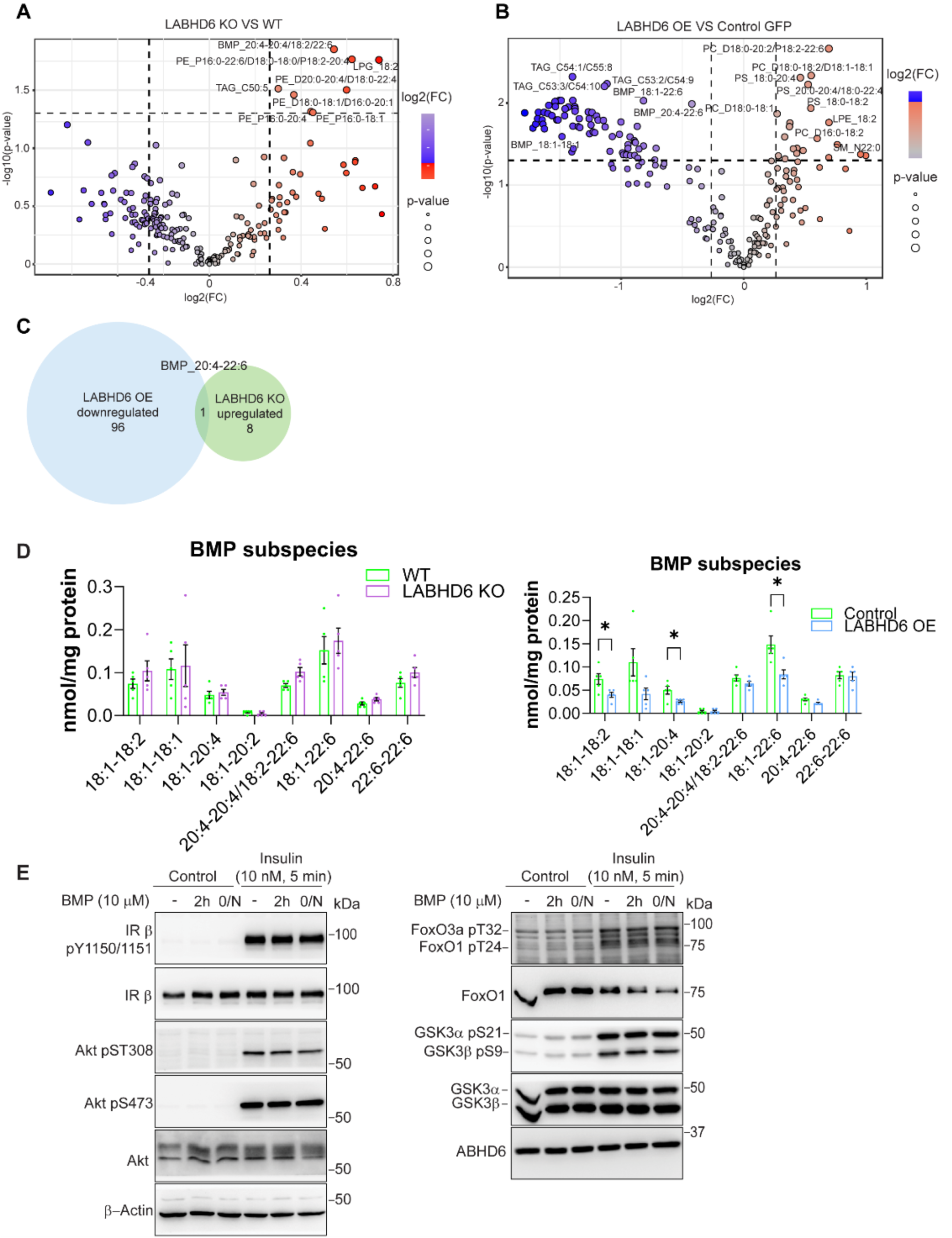
ABHD6 acts as a BMP hydrolase yet modulates insulin signaling independently of BMP. Livers from HFD-LABHD6 KO and HFD-LABHD6 OE mice (n = 5 per group) were used for lipidomic analysis. (A and B) Volcano plots of lipid species. (C) Venn diagram of differentially abundant lipid species. (D) Total BMP subspecies in livers of LABHD6 KO and OE mice. (E) Effects of exogenous BMP administration on hepatic insulin signaling. Data are displayed as mean ± SEM and analyzed by Student’s *t* test. *p < 0.05

**Sup Figure 5.**
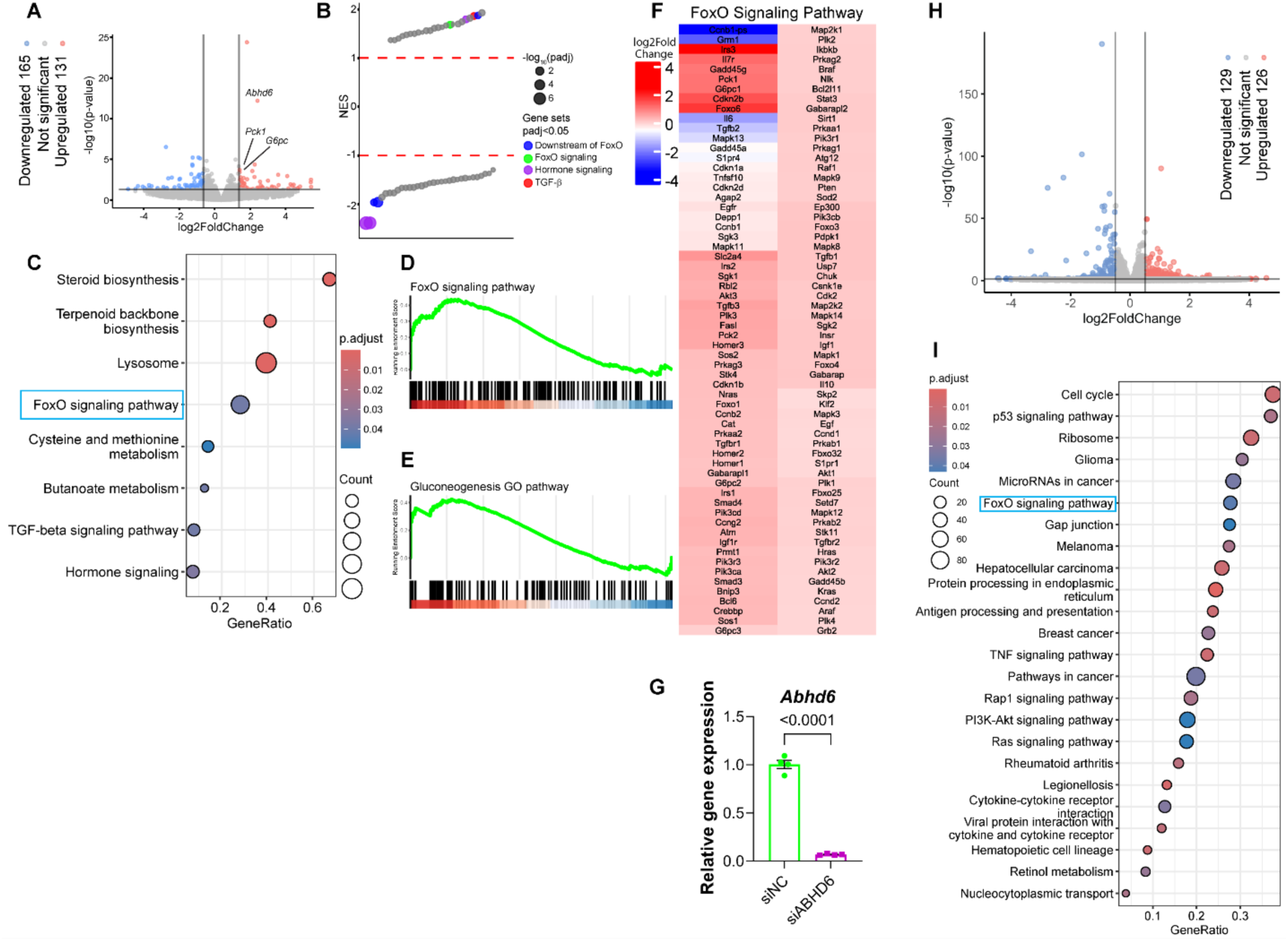
ABHD6 is a pivotal regulator of FoxO signaling. (A-F) 8-week-old male wildtype mice were injected with 1×10¹¹ genome copies AAV8 TBG-eGFP or TBG-*Abhd6* (n = 3 per group). 1 week post-injection, the mice were fed a high-fat diet (HFD) for 12 weeks. Bulk RNA sequencing was performed on livers. **(A)** Volcano plot of gene expression. **(B)** Upregulated and downregulated pathways. **(C)** Top ranked enriched pathways. **(D and E)** Gene Set Enrichment Analysis (GSEA) against ‘FoxO signaling’ and ‘Gluconeogenesis pathway’ sets. **(F)** Expression of genes in ‘FoxO signaling’ set. (G-I) Bulk RNA sequencing was performed in primary hepatocytes under conditions of ABHD6 knockdown by siRNA. (G) Verification of ABHD6 knockdown by siRNA. (H) Total changed genes in response to ABHD6 knockdown in primary hepatocytes. (I) Top ranked pathways that affected by ABHD6.

**Sup Figure 6.**
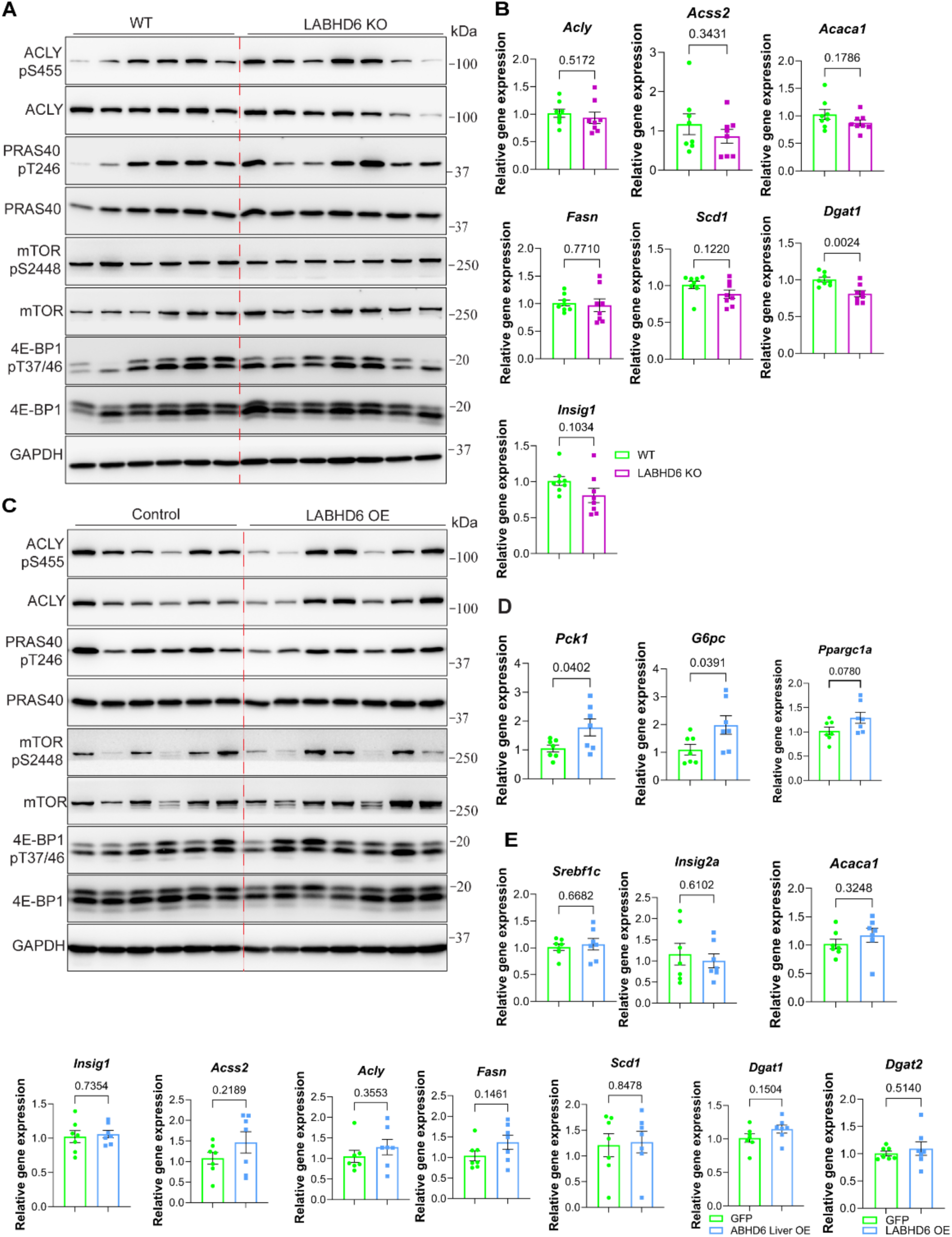
ABHD6 manipulation only affects Akt-Foxo1/3a-mediated gluconeogenesis, without affecting Akt-mediated lipogenesis. Liver samples were obtained from LABHD6 KO and OE mice after HFD feeding. Western blot and qPCR analysis were performed to check the hepatic insulin signaling. (A) Hepatic insulin signaling in LABHD6 KO mice. (B) Gene expression of Acly, Acss2, Acaca1, Dgat1, and Insig1. (C) Hepatic insulin signaling in LABHD6 OE mice. (D) Gene expression of gluconeogenic genes, including *Pck1*, *G6pc* and *Ppargc1a* in the livers of Control and LABHD6 OE mice. (E) Gene expression lipogenic genes, including *Srebf1*, *Insig2a*, *Acaca1*, *Insig1*, *Acss2*, *Acly*, *Fasn*, *Scd1*, *Dgat1* and *Dgat2*., . Data are displayed as mean ± SEM and analyzed by Student’s *t* test p-values of respective comparisons are provided.

**Sup Figure 7.**
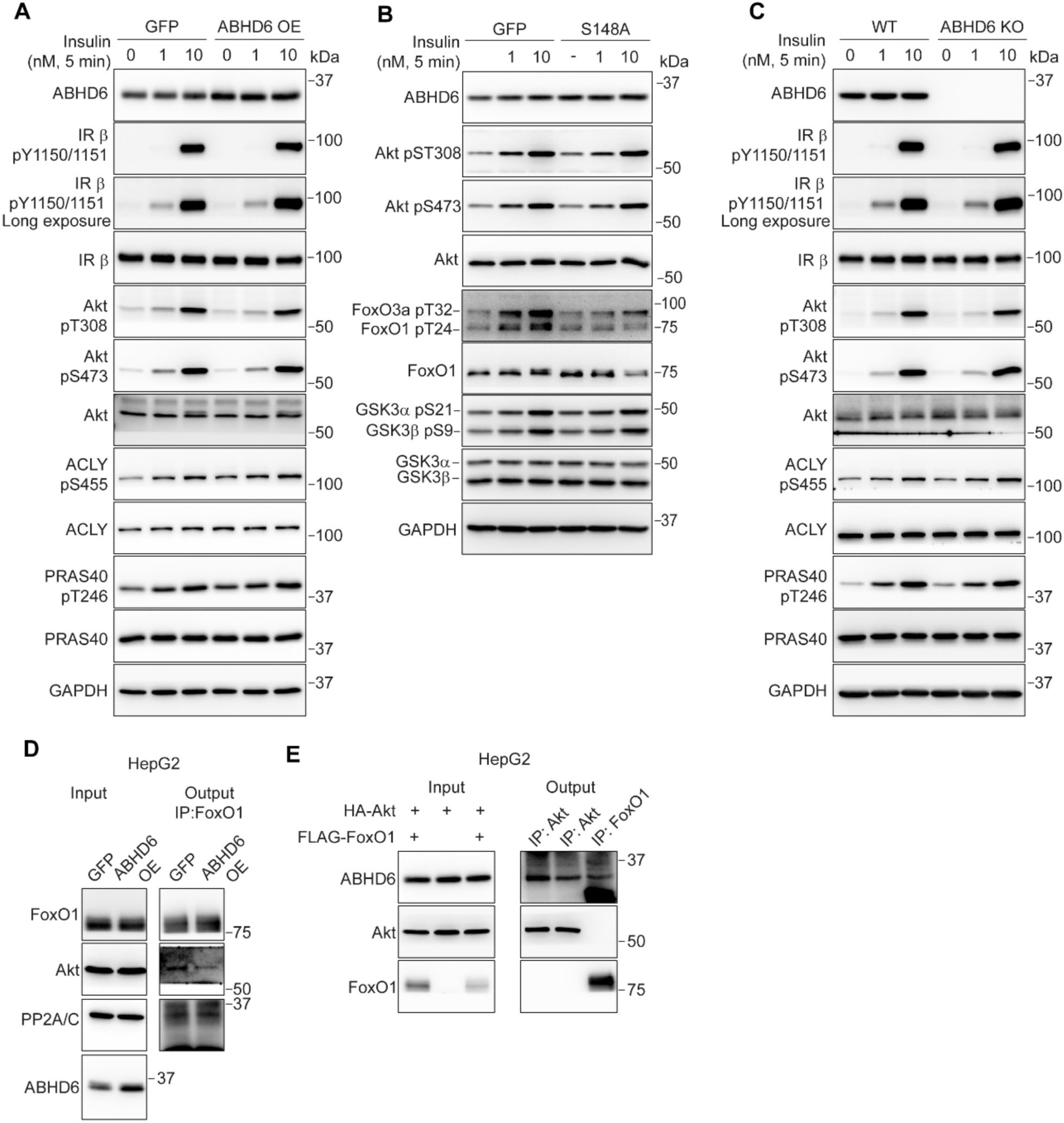
ABHD6 regulates Akt-Foxo1/3a phosphorylation and interaction, leading to altered gluconeogenesis. (A) Effects of ABHD6 overexpression on hepatic insulin signaling.(B) Effects of ABHD6 S148A overexpression on hepatic insulin signaling. (C) Effects of ABHD6 knockout on hepatic insulin signaling. (D) FoxO1 interaction with PP2A/C in ABHD6 OE HepG2 cells. HepG2 cells transduced with AAVDJ CAG-Abhd6 (ABHD6 OE) or CAG-eGFP (GFP) at MOI of 1000. 48 hours post-transfection/transduction, FoxO1 co-immunoprecipitation was performed. (E) ABHD6 interaction with Akt2 or FoxO1 in HepG2 cells. HepG2 cells were transfected with 1 μg HA-Akt2 and/or FLAG-FoxO1 plasmids and transduced with AAVDJ CAG-Abhd6 at MOI of 1000. 48 hours post-transfection/transduction, Akt2 and FoxO1 co-immunoprecipitation was performed.

**Sup Figure 8.**
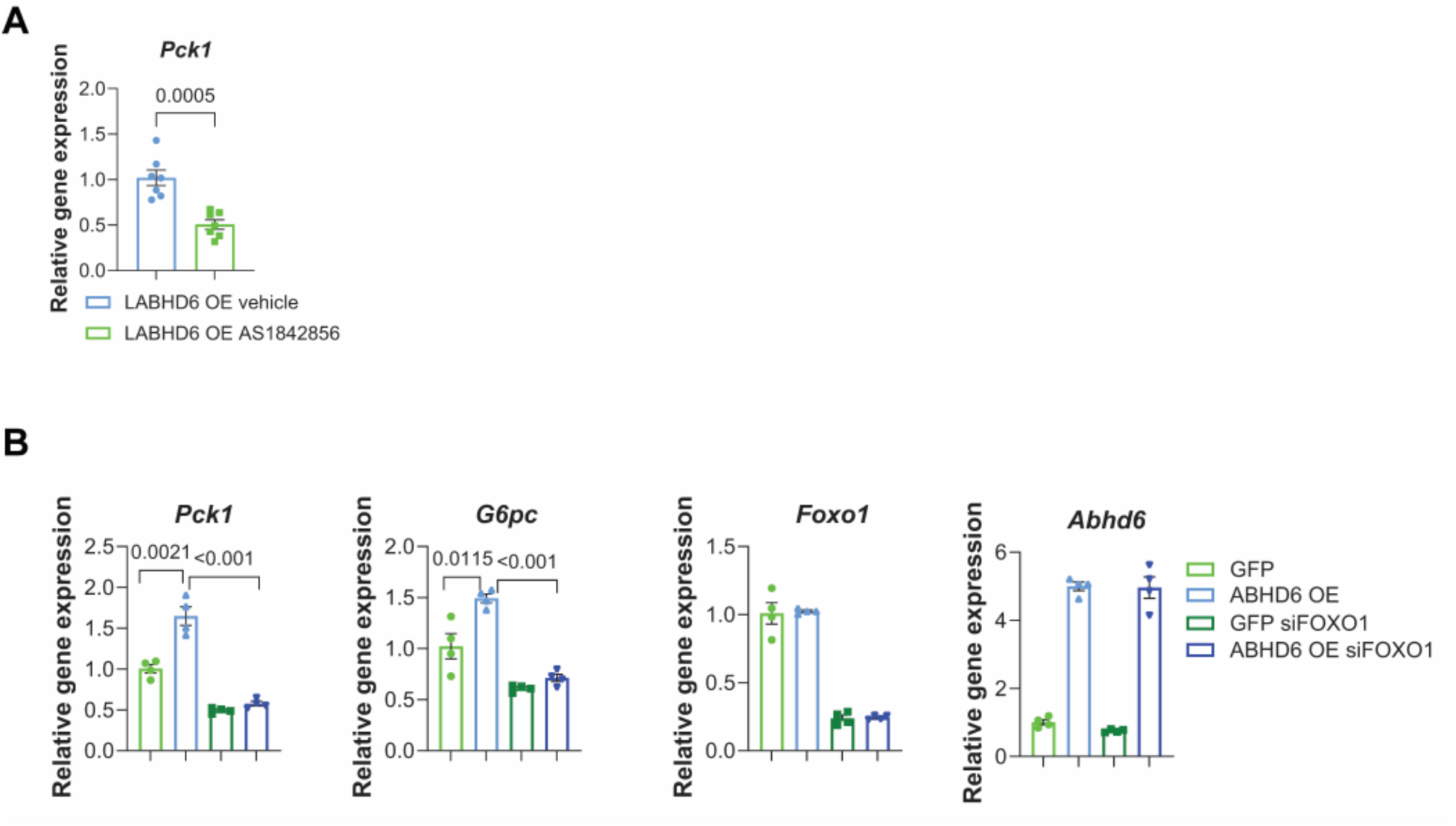
ABHD6 regulates insulin sensitivity in a FoxO1-dependent manner. (A) 8-week-old male wildtype mice were injected with 1×10¹¹ genome copies TBG- Abhd6 (LABHD6 OE). One week post-injection, the mice were fed a high-fat diet (HFD) for 12 weeks. AS1842856 was administered orally at 30 mg/kg, three times over two days (see Methods for experimental details). Gene expression of *Pck1* in the livers. (B) Foxo1 knockdown completely abolishes the effects of ABHD6 overexpression on gluconeogenic gene expression in primary hepatocyte. Data are displayed as mean ± SEM and analyzed by Student’s t test. p-values of respective comparisons are provided.

**Supplemental Table 1.**
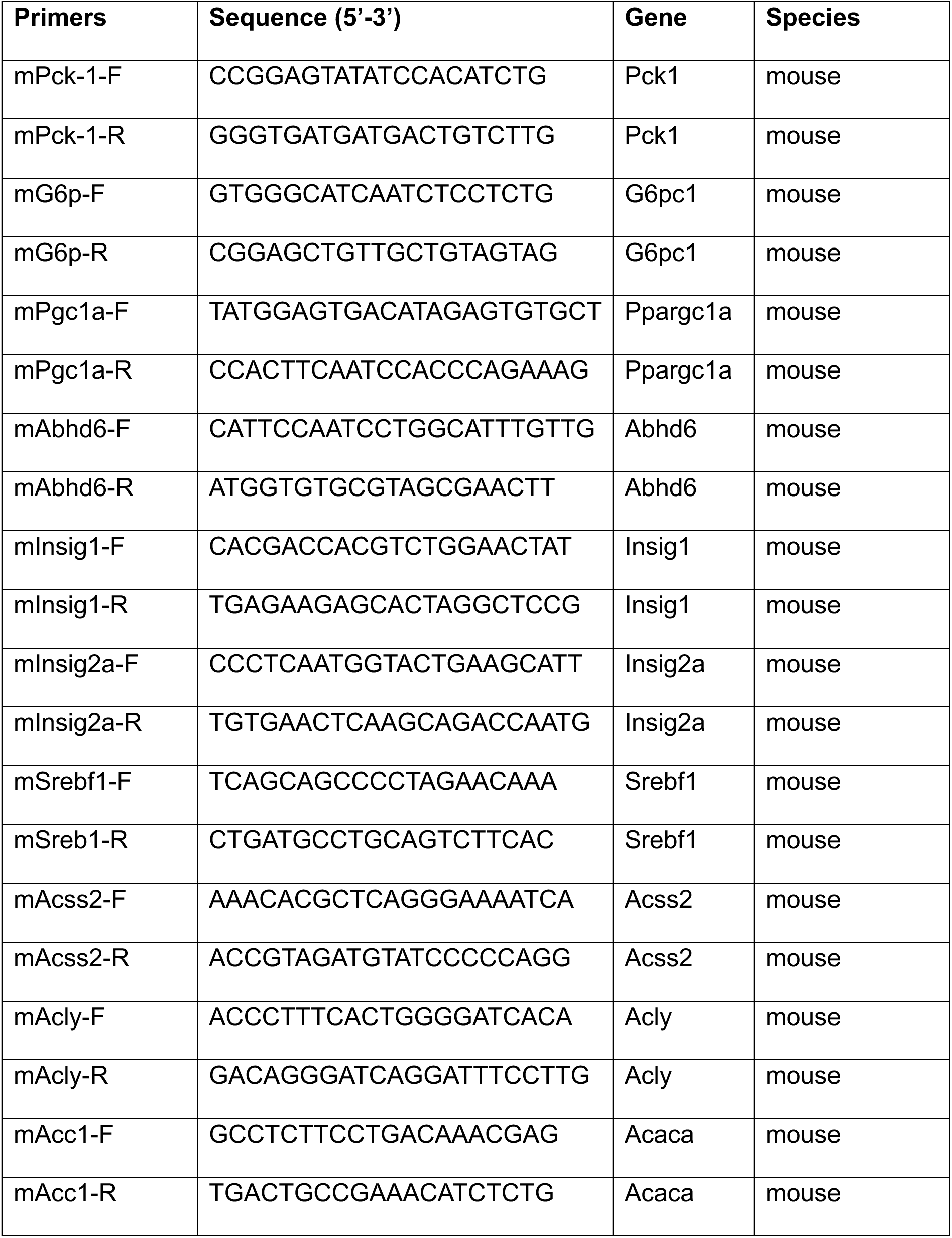
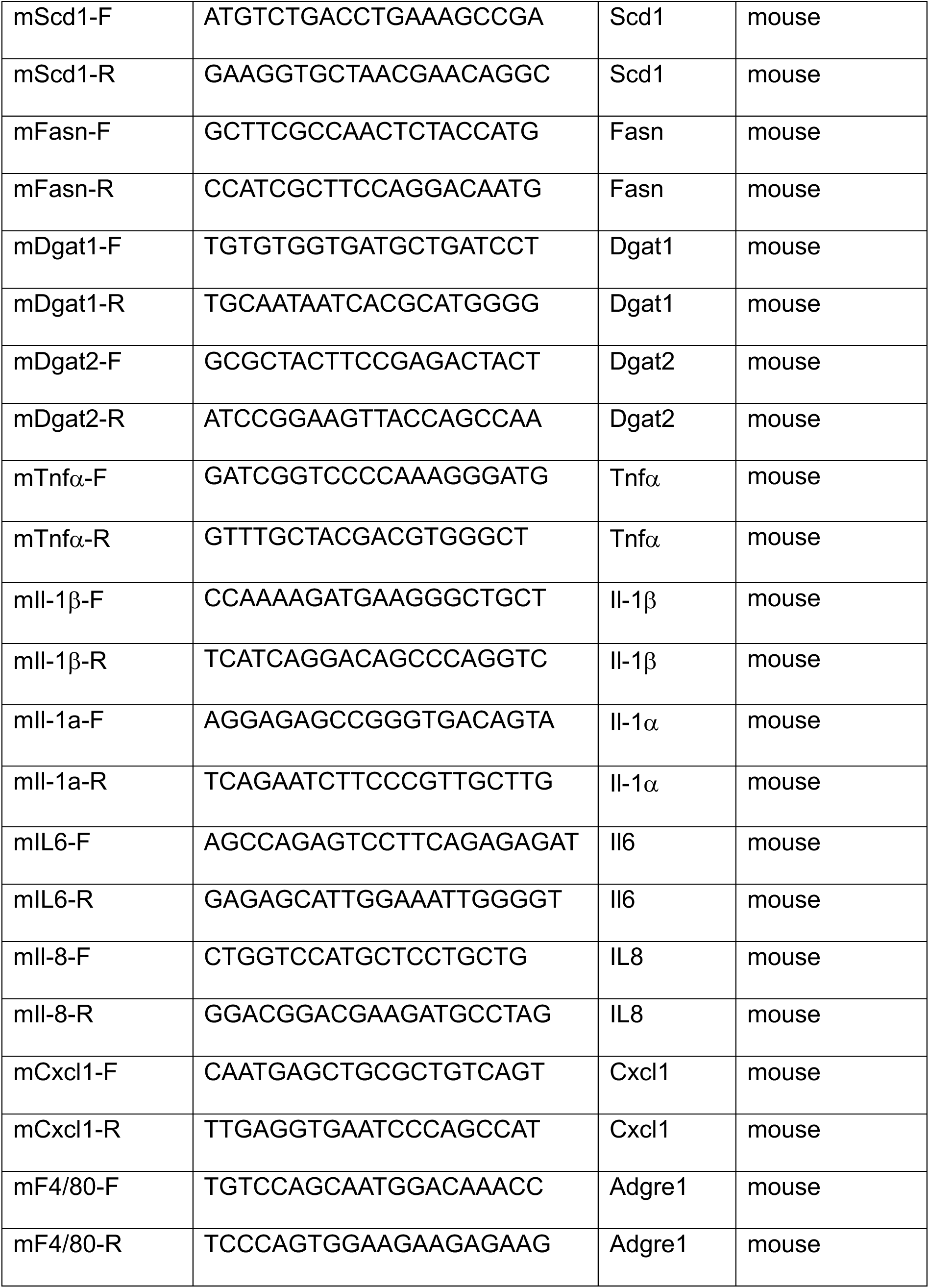
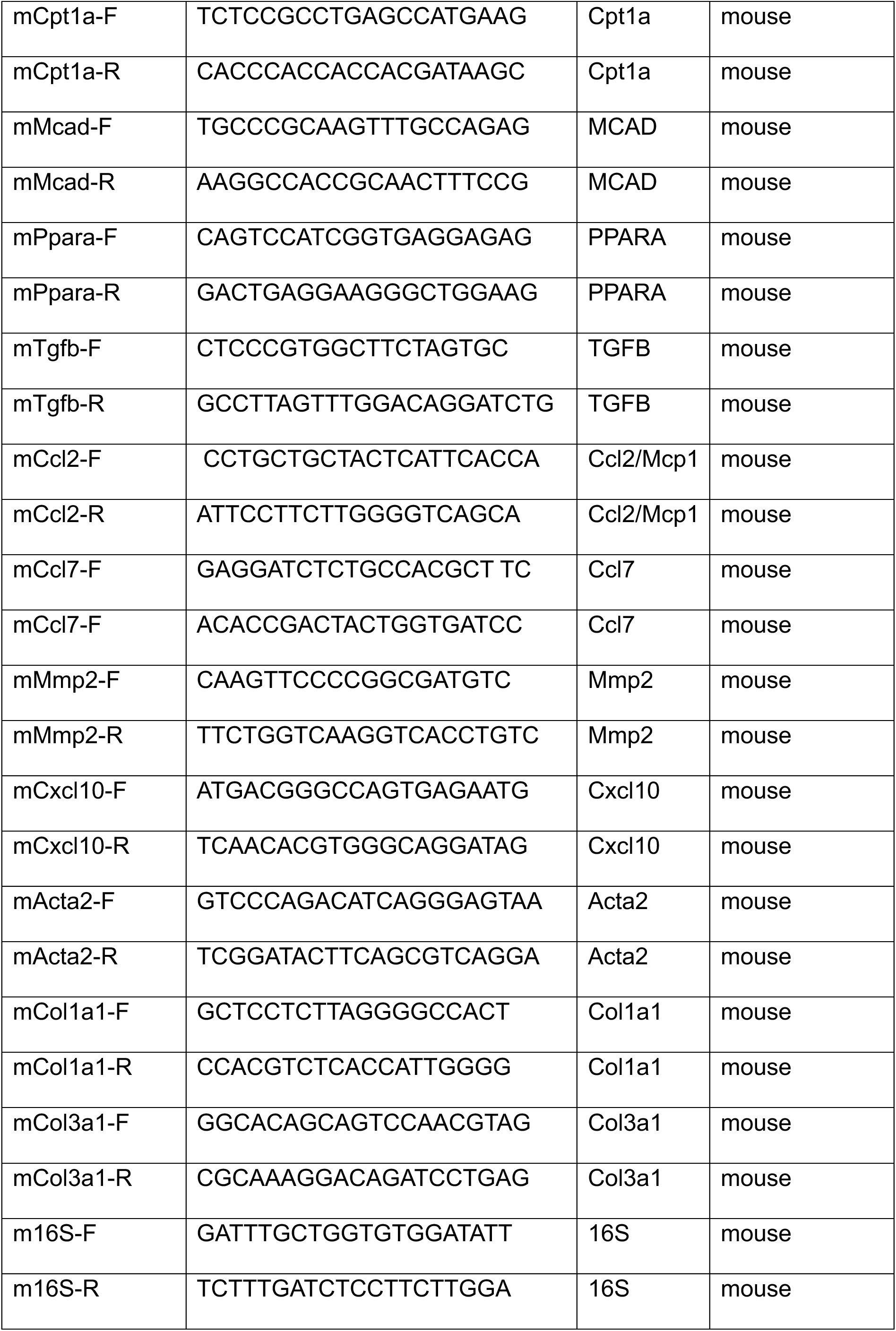
Primers used in the study.

**Supplemental Table 2.**
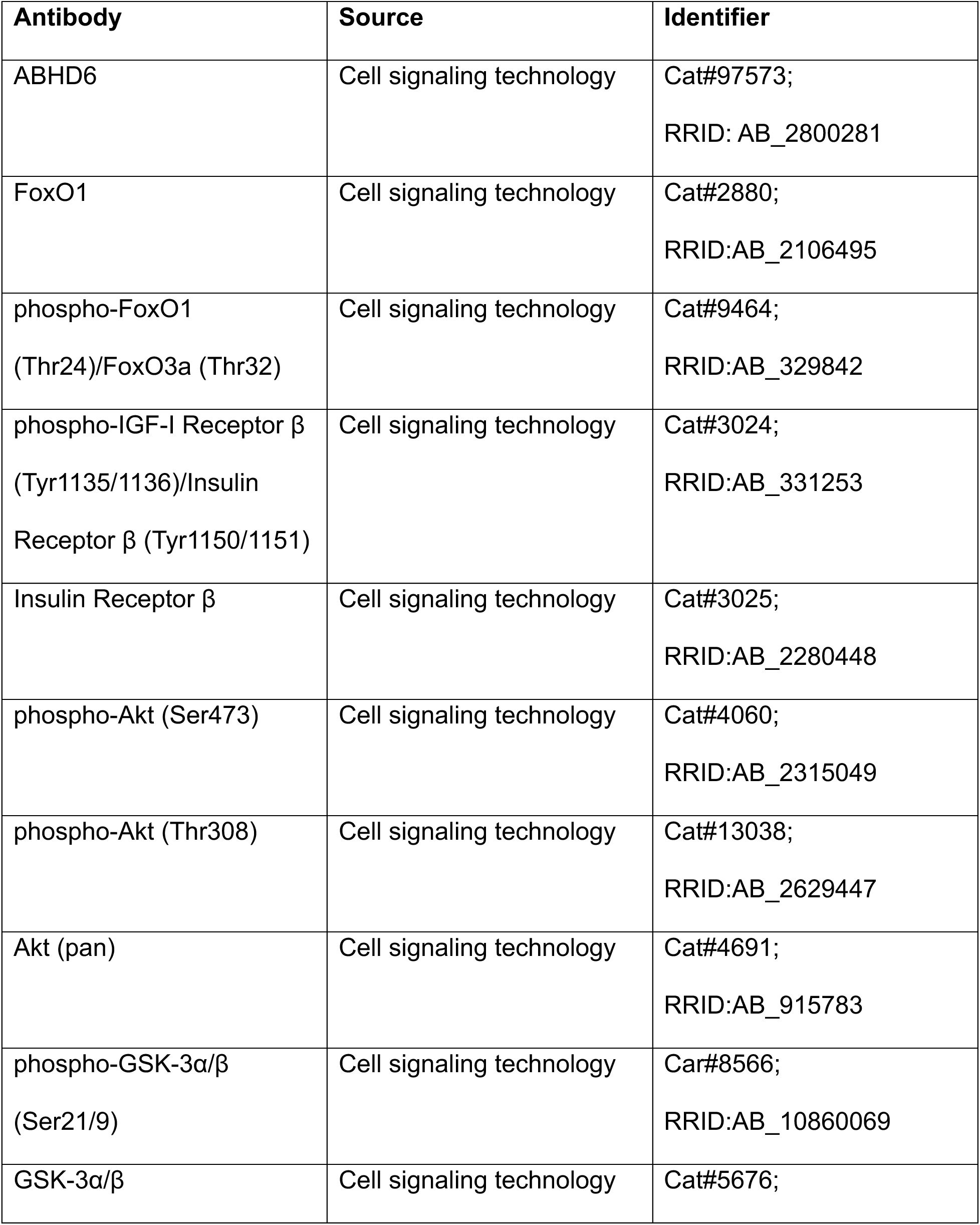
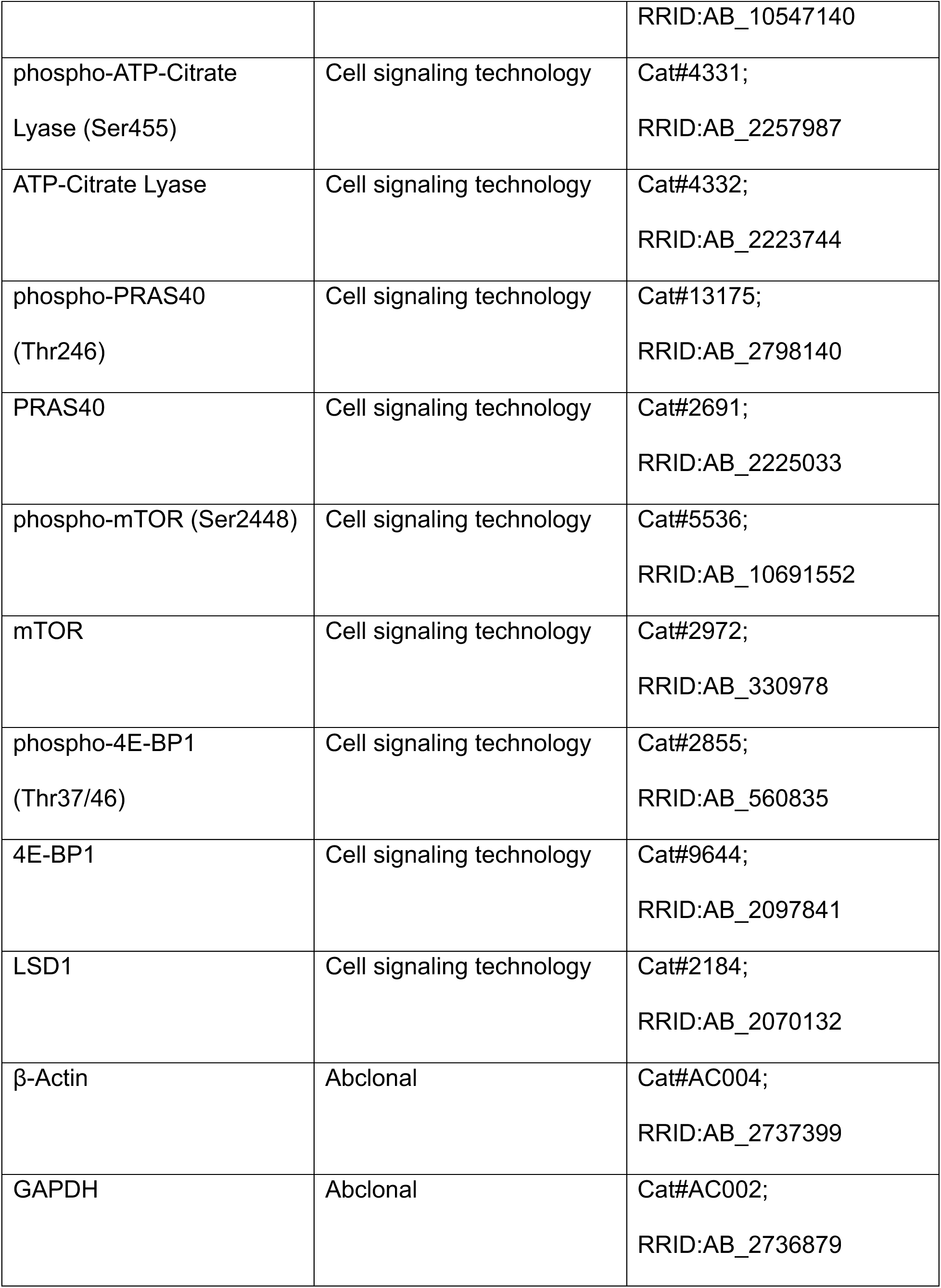
Antibodies used in the study.

## Notes

### Competing Interest Statement

The authors have declared no competing interest.

